# Deep learning-based brain transcriptomic signatures associated with the neuropathological and clinical severity of Alzheimer’s disease

**DOI:** 10.1101/2021.06.08.447615

**Authors:** Qi Wang, Kewei Chen, Yi Su, Eric M. Reiman, Joel T. Dudley, Benjamin Readhead

## Abstract

Brain tissue gene expression from donors with and without Alzheimer’s disease (AD) have been used to help inform the molecular changes associated with the development and potential treatment of this disorder. Here, we use a deep learning method to analyze RNA-seq data from 1,114 brain donors from the AMP-AD consortium to characterize post-mortem brain transcriptome signatures associated with amyloid-β plaque, tau neurofibrillary tangles, and clinical severity in multiple AD dementia populations. Starting from the cross-sectional data in the ROSMAP cohort (n = 634), a deep learning framework was built to obtain a trajectory that mirrors AD progression. A severity index (SI) was defined to quantitatively measure the progression based on the trajectory. Network analysis was then carried out to identify key gene (index gene) modules present in the model underlying the progression. Within this dataset, SIs were found to be very closely correlated with all AD neuropathology biomarkers (R ∼ 0.5, p < 1e-11) and global cognitive function (R = -0.68, p < 2.2e-16). We then applied the model to additional transcriptomic datasets from different brain regions (MAYO, n = 266; MSBB, n = 214), and observed that the model remained significantly predictive (p < 1e-3) of neuropathology and clinical severity. The index genes that significantly contributed to the model were integrated with AD co-expression regulatory networks, resolving four discrete gene modules that are implicated in vascular and metabolic dysfunction in different cell types respectively. Our work demonstrates the generalizability of this signature to frontal and temporal cortex measurements and additional brain donors with AD, other age-related neurological disorders and controls; and revealed the transcriptomic network modules contribute to neuropathological and clinical disease severity. This study illustrates the promise of using deep learning methods to analyze heterogeneous omics data and discover potentially targetable molecular networks that can inform the development, treatment and prevention of neurodegenerative diseases like AD.

## Introduction

As the age of the global population advances, dementia, with late-onset Alzheimer’s disease (LOAD) as the most prevalent form, has become a formidable public health threat. Despite numerous recent scientific advances in illuminating the pathophysiology of LOAD, no disease modifying treatments are currently available. This fact underscores the complicated molecular etiology driving the disease and the urgent need to broaden our search for effective therapeutics beyond the conventional amyloid cascade hypothesis.^1^

For a highly heterogeneous, multifactorial disease such as LOAD, integrated and large-scale genomic data analyses have been carried out to disentangle and capture the diverse gene regulatory interactions.^2^ Most of these studies focus on the exploration of the molecular mechanism of AD pathology by employing a case-control study design or modeling it as discrete stages, usually excluding the study subjects with other dementia pathologies, or omitting the mild cognitive impairment (MCI) stage and its role in the disease progression, even in the studies aiming to reveal the transcriptional dysregulation involving the progression of AD.^3^ Difficulties in sampling brain tissue throughout life coupled with globally limited access to diagnostic neuroimaging necessitates that a definitive diagnosis of AD is only made following postmortem neuropathological assessment. This further exacerbates the challenge we face in studying LOAD, or dementia in general as a continuous spectrum to find novel biomarker and drug targets. Recent efforts have begun to model AD progression as a continuous trajectory using cross-sectional transcriptomic data,^4, 5^ by leveraging the methods developed in single-cell genomics^6, 7^ and machine learning.^8^ Iturria-Medina et al.^4^ adopted an unsupervised machine learning algorithm applied to gene expression microarray data and discovered a contrastive trajectory in multiple cohorts respectively. The trajectory has been demonstrated to strongly predict neuropathological severity in AD in each dataset. Mukherjee et al.^5^ applied a manifold learning method to RNA-seq data to define ordering across samples based on gene expression similarity and estimate the disease pseudotime for each sample. Disease pseudotime was strongly correlated with the burden of Aβ, tau, and cognitive dementia within subjects with LOAD. Although these unsupervised machine learning methods have been shown to be highly predictive for well-known pathological biomarkers within a dataset, it would be desirable to have a generalized, universally predictive model for AD neuropathology and cognitive impairment across distinct cohorts and brain tissues, which helps decipher common AD etiology at molecular level. In addition, their broader application in peripheral tissues to identify novel biomarkers would greatly facilitate early diagnosis and progression monitoring of AD.

Deep learning methodologies are a rapidly evolving class of machine learning algorithms that have demonstrated superior performance over traditional machine learning approaches in identifying intricate structures in complex high-dimensional data, across diverse domains including computer vision, pattern recognition and bioinformatics.^9^ Specific to genomic data, it has been demonstrated that “big data” in many human diseases can be exploited by deep learning methods for early detection,^10^ disease classification,^11, 12^ and biomarker identification,^8, 13^ mostly in the cancer research field. More recent efforts have begun to apply similar methods towards research questions within the neuroscience research field,^14^ including the study of neurodegenerative disease,^15^ though the potential for these methods to contribute to novel insights in AD research remains underexplored.

In this work, we are leveraging the multi-dimensional, well characterized and high quality genomic, neuropathological and clinical data from the Accelerating Medicines Project for Alzheimer’s Disease (AMP-AD) program^16^ and applying the latest deep learning framework to identity pseudo-temporal trajectories in transcriptomic space and the underlying gene signatures for AD progression. As a major component of the AMP-AD program, the Target Discovery and Preclinical Validation Project brings together different organizations to collect and analyze multidimensional molecular data (genomic, transcriptomic, epigenomic, proteomic) from more than 2,000 human brains and peripheral tissues from multiple AD cohorts.^17^ Using the RNA-seq data from dorsolateral prefrontal cortex (DLPFC) region in the Religious Orders Study and Memory and Aging Project (ROSMAP) cohort,^18, 19^ we first trained a deep learning model to perform supervised classification between the two termini of the disease continuum (AD and control diagnosis group). The goal is to achieve the maximum separation of neuropathologically confirmed cases and controls. The model was subsequently applied to all the subjects within the cohort, and the intermediate layer of the obtained manifold for all subjects was further dimensionality reduced by Uniform Manifold Approximation and Projection (UMAP)^20^ to obtain a trajectory in three dimensional (3D) space for AD progression. We then derived an index to assess the stage of the progression, namely the severity index (SI) along the trajectory. We observed that the SI was significantly correlated with all the neuropathological biomarkers and achieved excellent model metrics aligned with global cognitive function score. When the deep learning model trained on the ROSMAP cohort was applied to two independent AMP-AD datasets, the MAYO RNA-seq study cohort^21^ and The Mount Sinai Brain Bank (MSBB) study cohort,^22^ similar trajectories and sample distribution following a generalized pattern were observed and the estimated SI values remain to be strongly correlated to pathological biomarkers and clinical severity. The model identified 593 genes (“index genes”) playing significant roles for the severity of AD-related neuropathology and cognitive impairment in the disease continuum. Network analysis suggests that these genes are clustered in six gene co-expression modules, four of which are strongly associated with neuropathology and clinical severity. One of the four modules shows especially high correlation with all the neuropathological biomarkers and clinical cognitive functions and the genes are associated with metabolic and vascular dysfunction in oligodendrocytes. The other three modules are also found to be associated with the pathological and clinical severity significantly in neurons, astrocytes, and endothelial cells respectively. Our results collectively demonstrate that a deep learning approach can reveal novel genomic information from complex, high dimensional gene expression data in a manner that can elucidate the molecular mechanisms of AD. The model can be readily applied to additional gene expression datasets to predict AD severity, thus indicating its potentially broad utility for AD diagnosis and staging. The approach also provides a general framework for studying multi-omics data to capture underlying molecular signatures towards novel biomarkers and drug targets of neurogenerative diseases.

## Materials and methods

### RNA-seq datasets from AMP-AD consortium

All the RNA-seq data were obtained from the AMP-AD data portal through Synapse (https://www.synapse.org/). Demographic information for each of the cohort (ROSMAP, MAYO and MSBB) sampled in the RNA-seq study is reported in supplementary Table S1. The processed, normalized data were obtained for each cohort respectively, from the harmonized, uniformly processed RNA-seq dataset across the three largest AMP-AD contributed studies (syn17115987). In ROSMAP cohort, all the brain tissue samples were collected from dorsolateral prefrontal cortex (DLPFC, n = 639, syn8456629).^18, 19^ In Mayo RNA-seq study (syn8466812), brain tissue samples were collected from cerebellum (CER, n = 275) and temporal cortex (TCX, n = 276).^21^ The MSBB study (syn8484987) has 1,096 samples from the Mount Sinai/JJ Peters VA Medical Center Brain Bank, which were sequenced from 315 subjects from four brain regions including frontal pole (FP, Brodmann area 10), inferior frontal gyrus (IFG, Brodmann area 44), superior temporal gyrus (STG, Brodmann area 22), and parahippocampal gyrus (PHG, Brodmann area 36) respectively.^22^ The harmonized processing of each study from three cohorts was previously performed using a consensus set of tools with only library type-specific parameters varying between pipelines (https://github.com/Sage-Bionetworks/ampad-DiffExp).^23^ The logCPM values from each dataset were used in all the subsequent analyses.

### Phenotypic data

All the clinical and pathological data for the ROSMAP cohort were obtained from the Rush Alzheimer’s Disease Center (RADC) Research Resource Sharing Hub (https://www.radc.rush.edu/home.htm), upon approval of data usage agreement. The following phenotypical measurements were used in the study: cogdx = final consensus cognitive diagnosis; age_death = age at death; educ = years of education; msex = sex; race7 = racial group; apoe4 = apoe4 allele count; PMI = postmortem interval; r_pd = clinical Parkinson’s disease; r_stroke = stroke diagnosis; dlbdx = pathologic diagnosis of Lewy body diseases; hspath_typ = hippocampal sclerosis; arteriol_scler = arteriolosclerosis; braaksc = Braak stage; ceradsc = CERAD score; gpath = global AD pathology burden; niareagansc = NIA-Reagan diagnosis of AD; amyloid = overall amyloid level; plaq_d = diffuse plaque burden; plaq_n = neuritic plaque burden; nft = neurofibrillary tangle burden; tangles = tangle density; cogn_global = global cognitive function. All the clinical diagnosis data were from the last visit, except for cogn_global, which was from the last available test. Their detailed definitions, together with possible values are reported in Supplementary Table S2. Among them, cogdx, braaksc and ceradsc values were used to define the class label for AD, control (CN) and OTHER groups (see below, methods for deep learning).

For MAYO and MSBB cohorts, subject clinical and pathological data were obtained from Synapse (syn3817650 for Mayo temporal cortex samples, syn5223705 for Mayo cerebellum samples, and syn6101474 for all the MSBB samples). For MAYO cohort, the following phenotypical data were used in the linear regression: age_death = age at death; gender = sex; apoe4 = apoe4 allele count; RIN = RNA integrity number; PMI = postmortem interval; Braak = Braak stage; Thal = Thal amyloid stage. For MSBB cohort, the following phenotypical data were used in the linear regression: age = age at death; sex = sex; race = racial group; apoe4 = apoe4 allele count; RIN = RNA integrity number; PMI = postmortem interval; Braak = Braak stage; PlaqueMean = mean plaque burden; CDR = clinical dementia rating; CERAD = CERAD score. The original CERAD score in the MSBB cohort was defined as: 1=Normal, 2=Definite AD, 3=Probable AD, 4=Possible AD. They were recoded to be semiquantitative as follows: 1=Definite AD, 2=Probable AD, 3=Possible AD, and 4=Normal, to be consistent with the notion used in the ROSMAP cohort.

### Deep learning of the transcriptome from DLPFC tissues in ROSMAP cohort

The whole machine learning framework consists of two major components, supervised classification (deep learning) and unsupervised dimension reduction. The deep learning method was built wholly based on the approach implemented in the previous implementation DeepType (https://github.com/runpuchen/DeepType).^11^ The detailed algorithm could be found in the reference. The method has been demonstrated to achieve superior performance on independent datasets and is very robust against label noise in classifying genomic data from complex human diseases such as cancer.^24^ In this work, we incorporated the method into our model (supervised classification) and applied it to the normalized logCPM data from ROSMAP cohort, which consists of the expression profile of 634 subjects with various AD pathology for 15,582 genes, and further applied an unsupervised dimension reduction method to obtain the pseudo-temporary trajectory for AD progression. The whole framework is illustrated in Figure 1.

**Figure 1.**
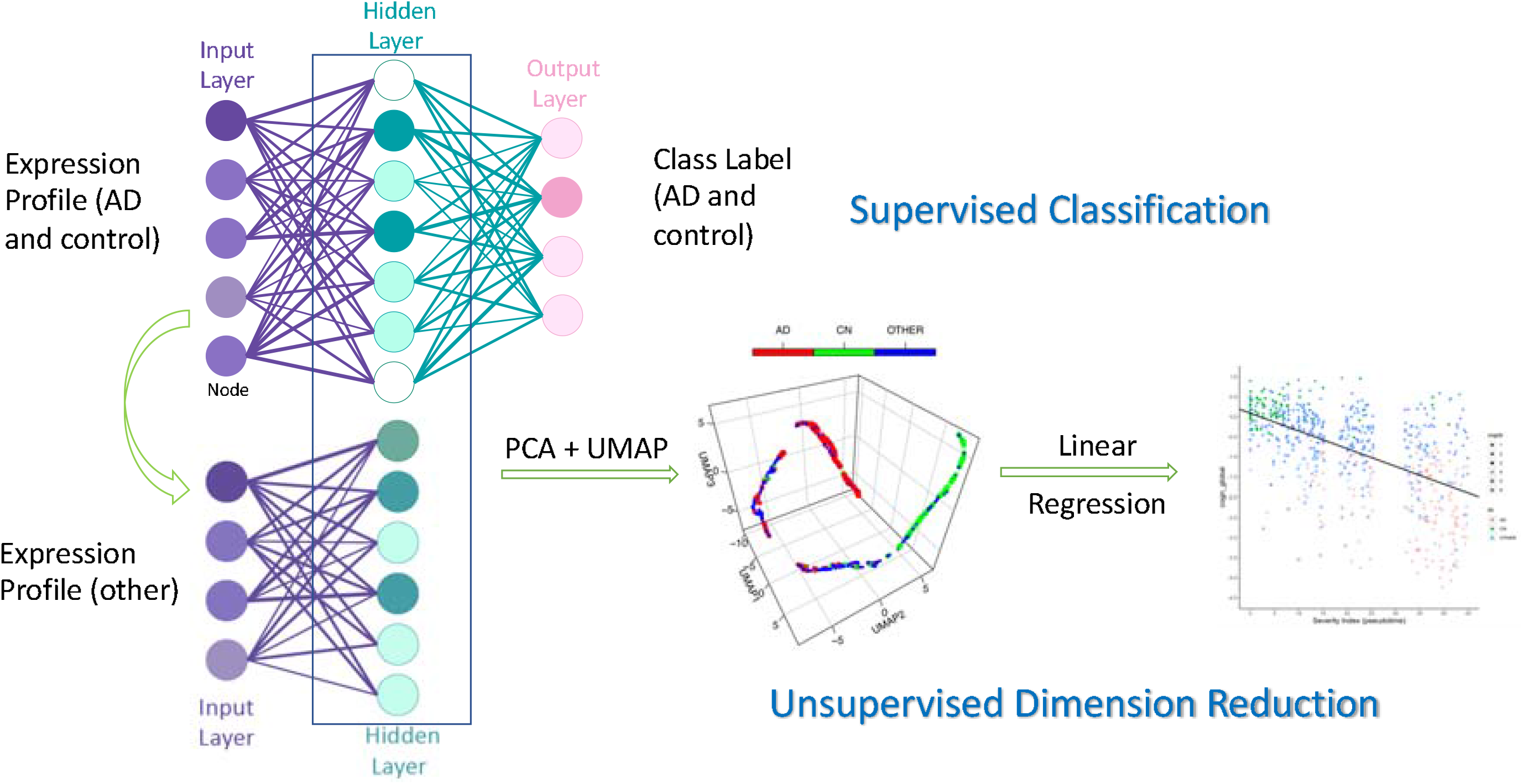
The deep learning framework employed in this work. Using the gene expression profiles from AD and control subjects and their diagnosis class as the input for supervised classification, the model was trained by a three-layer neural network. Then the response function from input layer to the hidden layer was applied to the profiles from the whole cohort. The resulting manifold was subject to unsupervised dimension reduction (PCA + UMAP) to obtain the pseudo-temporal trajectory and SI. SI was linearly correlated with phenotypic data for evaluation.

For the deep learning step, we used neuropathologically confirmed AD patients and normal controls, the two termini of the AD continuum, to train the model and identify transcriptomics signatures that differentiate the two groups. Interpretation of the diagnosis was as following:

> AD (156 samples): cogdx = 4, braaksc >= 4 and ceradsc <= 2;
>
> CN (control, 87 samples): cogdx = 1, braaksc <= 3 and ceradsc >= 3;
>
> OTHER (391 samples): All the other samples.

This was consistent with the criteria used in the previous differential expression analysis (syn8456629).^23^ Genes were first sorted in a descending order by variance of logCPM values for the whole dataset. The deep learning model was first built for the 243 samples from AD and control diagnosis groups. Data were randomly partitioned into training and test datasets, containing 80% (195) and 20% (48) of the samples with balanced distribution from each group. The logCPM values in the training set were first converted to Z score, followed by scaling those in the test set to the same scale. A three-layer neural network was trained, with the number of the nodes in the input layer, the intermediate layer, and the output layer set to 15,582, 128, and 1, respectively. In DeepType, the Adam method^25^ was employed to tune the parameters of the model, and a semi-supervised approach was adopted to train three hyper-parameters: the number of clusters *K*, the trade-off parameter α and the regularization parameter λ. The learning rate was set to 1e-4, the number of training epochs for model initialization and the joint supervised and unsupervised training were set to 1,500 and 5,000, respectively, and the batch size was set to 256. The model was trained by 5-fold internal cross-validation for the training set and the optimal *K*, α and λ were determined by the cross-validation to be 2, 2 and 0.004, respectively. Training and validation losses in the training process were tracked to avoid over-fitting.

After the training process was accomplished, a manifold representation of the intermediate layer was obtained for all the 634 samples in the whole cohort by forward pass using the trained network. Prior to that, data was scaled to the Z score using the same mean and standard deviation (SD) as the training set. The equation, as implemented in DeepType in MATLAB language, is as follows (eq 1-3):

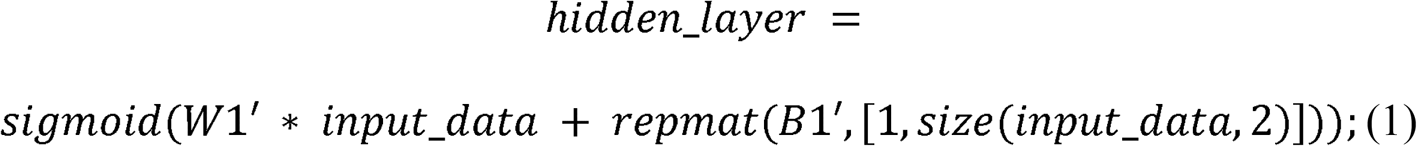

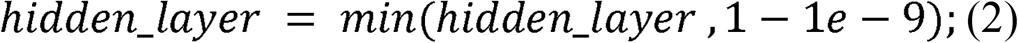

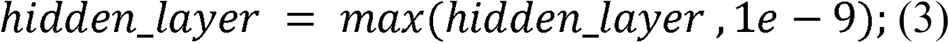

where *W1* is the first-layer weighting matrix and *B1* the first-layer bias vector obtained from the model. The hidden layer was bounded between (0, 1). Input_data was the expression matrix with data scaled and sorted in the same order as in the training set.

The resulting representation of the hidden_layer was further dimensionality reduced, first to 50 dimensions, by efficient computation of a truncated principal components analysis (PCA) using an implicitly restarted Lanczos method as implemented in the R package Monocle3,^26^ using the function preprocess_cds, without normalization or scaling. It was further reduced to the first three dimensions, by the Uniform Manifold Approximation and Projection (UMAP)^20^ method as implemented in the R package uwot (https://github.com/jlmelville/uwot). To ensure reproducibility, the following parameters were set: n_components = 3, nn_method=“annoy”, n_neighbors = 15L, metric = “cosine”, min_dist = 0.1, fast_sgd = F, ret_model = T, with random seed set to 2016.

### Severity index calculation and correlation with phenotypic data in ROSMAP cohort

Severity index (SI) for AD progression was derived for each sample, based on the 3D UMAP trajectory obtained earlier, by applying the method of inferring pseudotimes for single-cell transcriptomics from the function “slingPseudotime” as implemented in the R package Slingshot.^27^ SIs were then linearly correlated with all the AD clinical and pathological biomarkers individually, including the covariates r_pd, r_stroke, dlbdx, hspath_typ, arteriol_scler, PMI, RIN, apoe4, age_death, educ, msex, and race7 (detailed definitions can be found in Table S2, and data collection is reported in^28^), using the following linear regression model:

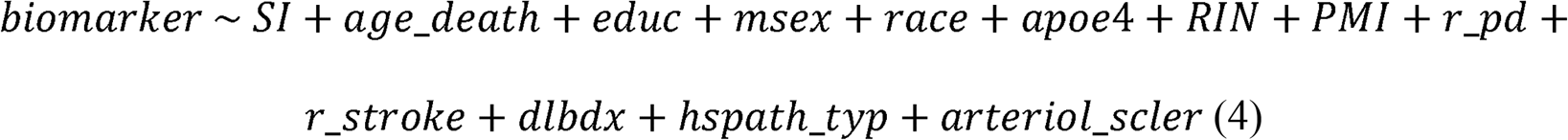

The pathological biomarkers and AD clinical measures used as dependent variables in the model are braaksc, ceradsc, niareagansc, gpath, amyloid, plaq_d, plaq_n, nft, tangles, and cogn_global. All the neuropathological measurements (gpath, amyloid, plaq_d, plaq_n, nft, tangles) were log transformed in the correlation analysis. All the semi-quantitative and quantitative measurements were treated as numerical; diagnosis of Lewy Body diseases, gender, and race were treated as categorical. Correlation coefficients were obtained by the “lm” function in R. Proportion of variance explained (PVE) for each predictor was obtained from the incremental sums of squares table by the “anova” function in R on the model, using the above order.

### Applying the deep learning model to external datasets (MAYO, MSBB)

The harmonized, uniformly processed RNA-seq datasets were first sorted by the same gene order as the input dataset of ROSMAP. Batch effects were then removed by the ComBat function^29^ in the R package sva.^30^ The input expression matrix subsequently was transformed to Z score by scaling to the training set in the deep learning model. A manifold representation was obtained for all the samples in each cohort by forward pass of the trained network, using eq 1-3) and reduced again to 50 dimensions by PCA. Trajectories were obtained by carrying out the UMAP transformation of the existing embedding model from ROSMAP DLPFC data, by the “umap_transform” function in R package uwot. SI for each sample was again derived from “slingPseudotime” function in Slingshot.^27^ Linear correlation of the SIs with all the pathological and clinical biomarkers were carried out by the “lm” function in R, using other non-AD pathology related variables as covariates when available (age, sex, race, PMI, RIN, apoe4 allele counts), by the following linear regression model:

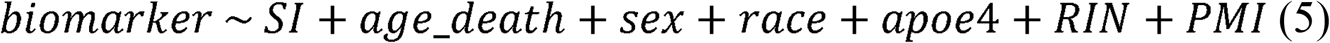

### Network and cell type analysis of the significant genes underlying AD progression

The hidden layer of the deep learning model returned a weight vector for each of the 15,582 genes in the input dataset. The root sum squares (RSS) of the weight vector for each gene was calculated, normalized to the maximal RSS and taken as the weight for each gene in the deep learning model. The weights from all genes were put into histogram in logarithm scale and a cut was made to separate the bimodal distribution. The genes in the higher weight groups were identified as significant genes contributing to AD progression (index genes).

Unsigned co-expression networks were built for these genes’ expression profile using the unscaled logCPM values. Network modules were identified using the cutreeDynamic function in the R package WGCNA,^31^ setting the minimum module size to 30. The power of 4 was chosen using the scale-free topology criterion. Correlation of 0.35, or height cut of 0.35 with deepSplit = 4 was used to merge modules whose genes are highly co-expressed.

Functional enrichment analysis was performed using Metascape,^32^ which uses a hypergeometric test and Benjamini-Hochberg p value correction to identify ontology terms that contain a statistically greater number of genes in common with an input list than expected by chance, using the whole transcriptome as background. Statistically significant enriched terms based on Gene Ontology,^33^ KEGG,^34^ Reactome,^35^ MSigDB^36^ were clustered based on Kappa-statistical similarities among their gene memberships. A 0.3 kappa score was applied as a threshold to identify enriched terms.

Fisher’s exact test was used to test enrichment of the gene set from each module with the gene sets generated for the ROSMAP samples from the meta-analysis of AD co-expression modules,^23^ or other curated AD gene sets (supplemental materials). Resulting p values were corrected using Bonferroni method for multiple test correction. Cell type enrichments were also done using Fisher’s exact test of gene set overlap with cell type specific gene sets from human reference single-cell RNA-seq data^37^ and the unique marker genes present in the single-cell RNA-seq data (with log2FC > 1) from the prefrontal cortical samples of AD patients and normal control subjects.^38^

Cell type marker gene expression signatures along SI were obtained by first smoothing each gene’s expression as a function of SI using a smoothing spline of degree of freedom = 3. The weighted mean of the marker genes was obtained and normalized to lie in [0,1]. The smoothed and normalized expression of marker genes for each cell type was plotted as a function of SI.

### Data availability

All the datasets from AMP-AD consortium used in this study are available at the AD knowledge portal (https://adknowledgeportal.synapse.org/), with synapse identifiers provided in the text. The machine learning framework, including the trained model, SIs for each cohort, and the codes to apply the trained neural network, map to the DLPFC 3D UMAP space, and obtain the SI will be uploaded to synapse upon publication of this work. The source code with synapse data withheld is available at https://github.com/qwang178/DeepBrain.

## Results

### Deep learning identified a pseudo-temporal trajectory for AD progression

We designed a three-layer deep learning model to dissect the gene expression data from DLPFC tissues across the AD spectrum. This simple scheme consists of inputting the two termini of the spectrum (i.e. pathologically confirmed AD and control groups) to obtain a learned representation encoded by the intermediate layer. We set the number of output clusters *K* at 2 throughout the learning process (i.e. we are not interested in any subcluster within the two termini). The regularization parameter λ and the trade-off parameter α were estimated to be 0.004 and 2, respectively (Supplementary Figure S1a, b). The training and validation losses in the training process were tracked and no sign of over-fitting was observed (Supplementary Figure S1c). The training and validation accuracy was observed to be stable at ∼97% and ∼90% respectively (Supplementary Figure S1d).

After the intermediate layer was mapped into 3D UMAP space, a prominent progressive trajectory, with two distinct clusters at both termini could be observed (Figure 2a). Mapping of the other samples into the same space clearly indicated a continuous disease spectrum as well as a progression course along the trajectory (Figure 2b). SI was calculated as the traveling distance along the trajectory by setting the starting point at the control terminus to zero, which reflects the disease progression of the subjects. When correlating with pathological biomarkers, the SI showed strong correlations with all the measurements (p <= 3.2e-6), with the weakest correlation observed for diffuse plaque, which was still highly significant (p = 3.2e-6) (Figure 2c). In addition, it indicated that APOE4 allele counts also contributed to all the biomarkers with various degrees of significance (p = 1.24e-04 to 1.47e­09), confirming it as a major genetic risk determinant for AD. Most strikingly, the model explained the greatest amount of variance for global cognitive function (R = -0.68) (Figure 2d), with SI contributing to the largest proportion of variance explained (PVE = 0.35, p < 2e­16, table S3). It also indicated that global cognitive function in this cohort was positively correlated with education (PVE = 0.0020, p = 5.57e-3), inversely correlated with APOE4 allele count (PVE = 0.035, p = 5.00e-6), a diagnosis of Parkinson’s disease (PVE = 0.042, p = 7.49e-7), neocortical Lewy Body disease (PVE = 0.016, p = 5.49e-4), hippocampal sclerosis (PVE = 0.014, p = 1.03e-3), and marginally age (PVE = 0.0047, p = 9.89e-2). When the two termini which were used in the training process were excluded from linear regression model, we still observed strong correlations between SI and all neuropathology biomarkers and clinical severity (p < 0.1), especially for global cognitive function (PVE = 0.15, p = 3.5e-7 for SI, R = -0.55 for the model, Table S3, S4, Figure S4), demonstrating the generality of the model outside the training data.

**Figure 2.**
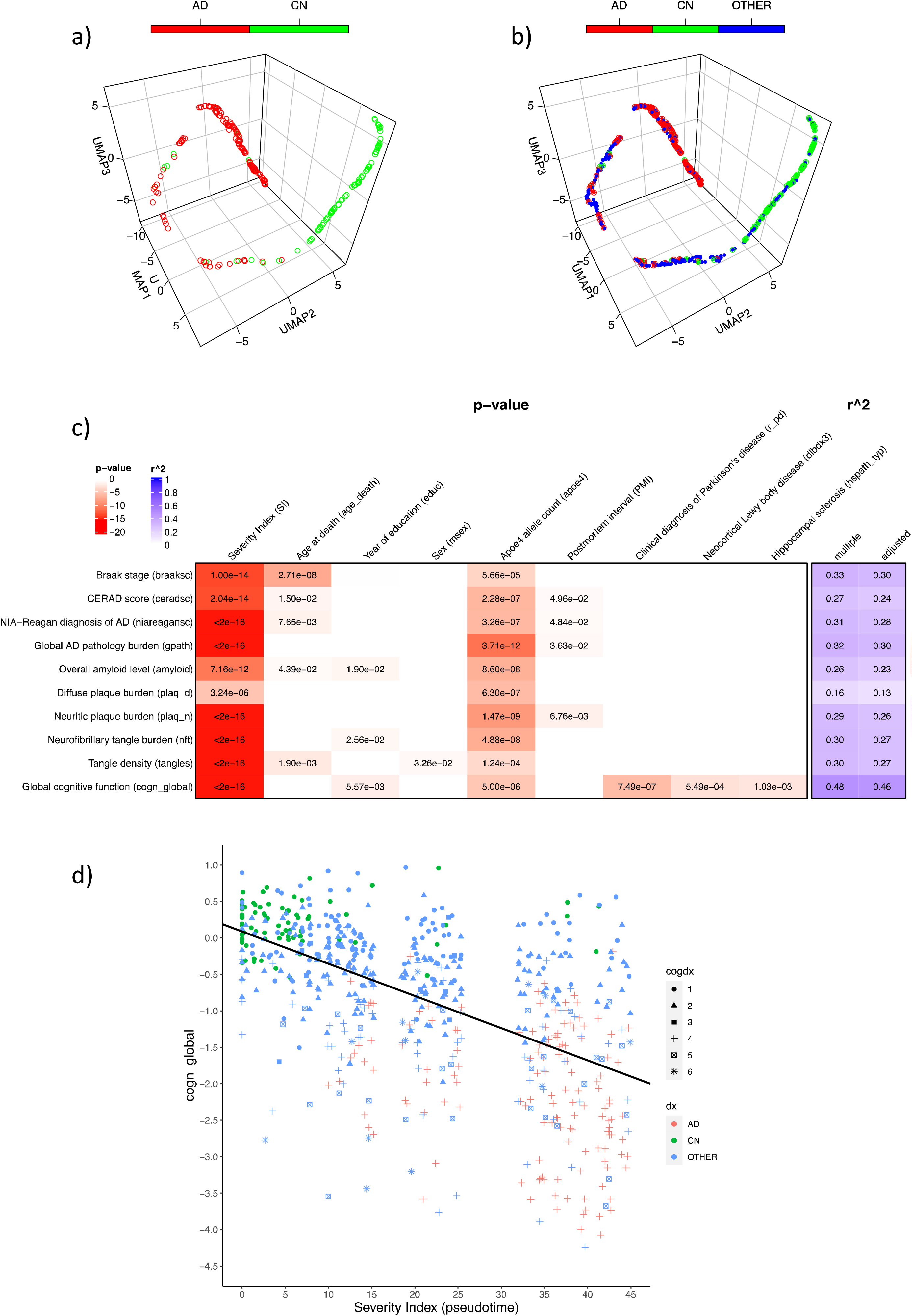
The pseudo-temporal trajectory from the trained deep learning model for the transcriptome from DLPFC tissues of ROSMAP cohort and the SI correlation with phenotypical data. Diagnosis class is defined in the main text. a) The trajectory with only AD and control (deep learning dataset) shown. b) The trajectory with all subjects (AD + CN + OTHER) shown. c) The model metrics for the linear regression between SI, controlled covariates and all the neuropathological biomarkers and global cognitive function, with p values shown in cells with p < 0.05. Only covariates with significant association with at least one biomarker were shown. The descriptions for the definition of each parameter can be found in Table S2. Detailed model metrics stratified by diagnosis groups can be found in Table S3 and S4. d) The linear regression plot between SI and global cognitive function. Color legend for cogdx (final consensus cognitive diagnosis):1= No cognitive impairment (CI); 2 = MCI and no other cause of CI; 3 = MCI and another cause of CI; 4 = AD and NO other cause of CI; 5 = AD and another cause of CI; 6 = Other dementia.

### Model achieved comparably strong performance in MAYO/MSBB cohorts

The model was applied to the harmonized transcriptomic data from both the MAYO and MSBB cohorts. Data from the MAYO cohort came from two different brain regions: temporal cortex (TCX) and cerebellum (CER). After projecting into the same 3D UMAP space, the subject distributions along the trajectories in the two different brain regions showed different patterns (Figure 3a, b). For TCX, it showed the distributions of different locations for AD vs control subjects along the trajectory similar to those from ROSMAP data, while this was not observed for CER as one would expect. It was also confirmed by the results obtained from linear regression of the SI vs pathological biomarkers (Braak and Thal scores, Figure 3c). Only in the TCX samples were the SIs found to be significantly correlated with both Braak (p = 4.88e-5) and Thal scores (p = 1.56e-3). Again the model explained a large amount of variance overall for both biomarkers, with R = 0.68. For MSBB cohort, the same model was applied to the gene expression profile of all four sampled regions (FP (BM10), STG (BM22), PHG (BM36), and IFG (BM44)), and all regions show similar albeit slightly different trajectories, with the SI consistently significantly correlated with all the neuropathological and clinical biomarkers (Braak score, PlaqueMean, CDR scale, and CERAD score, Figure 4).

**Figure 3.**
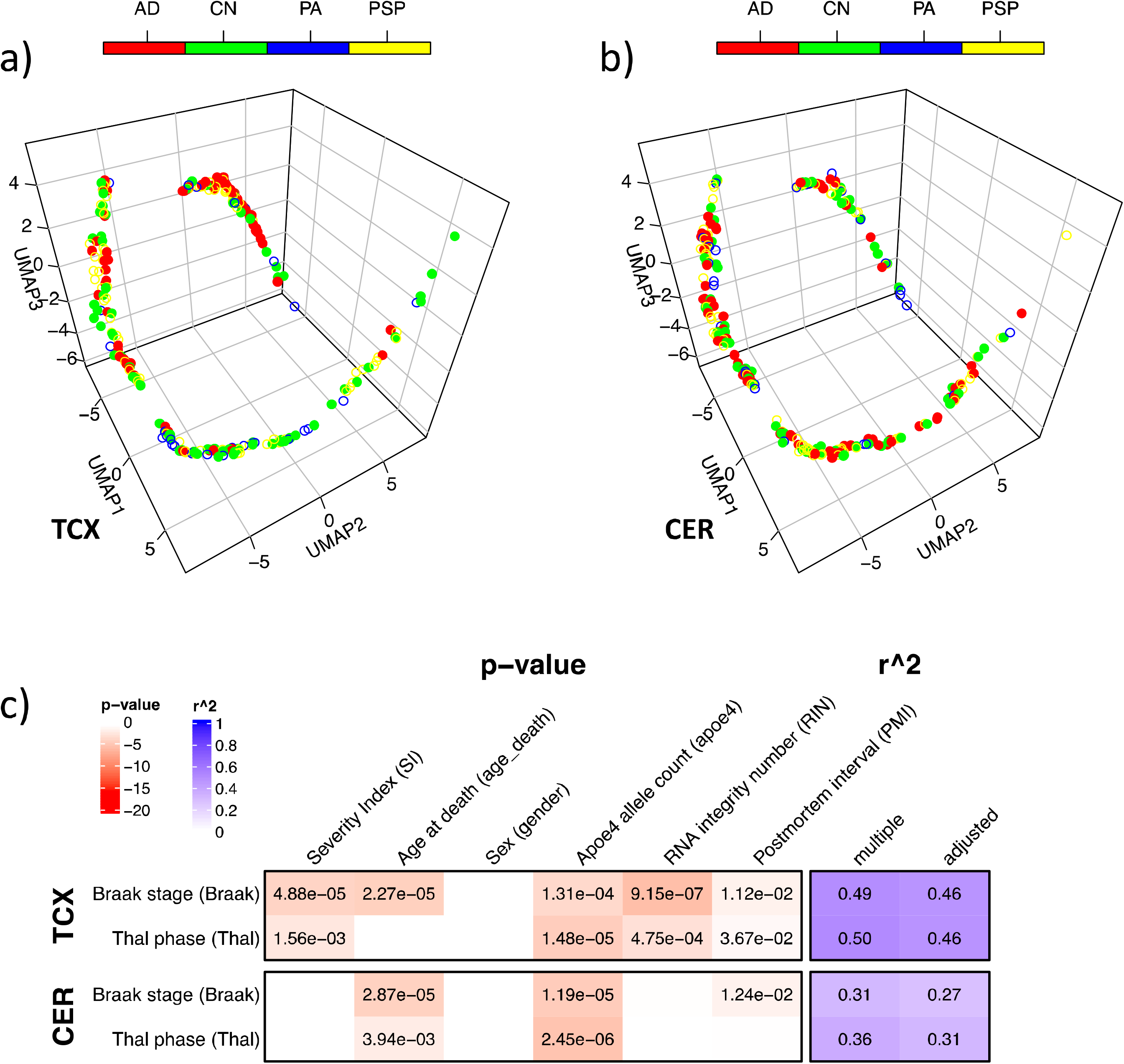
The pseudo-temporal trajectory for the transcriptomes from two brain regions of MAYO cohort and the SI correlation with phenotypic data generated by applying the trained deep learning model and mapping to the same 3D space as ROSMAP. AD = Alzheimer’s disease; CN = control; PA = pathological aging; PSP = progressive supranuclear palsy. a) The trajectory for transcriptome from TCX. b) The trajectory for transcriptome from CER. c) The model metrics for the linear regression between SI and the neuropathological biomarkers by different brain regions, with p values shown in cells with p < 0.05. Detailed model metrics are reported in Table S5.

**Figure 4.**
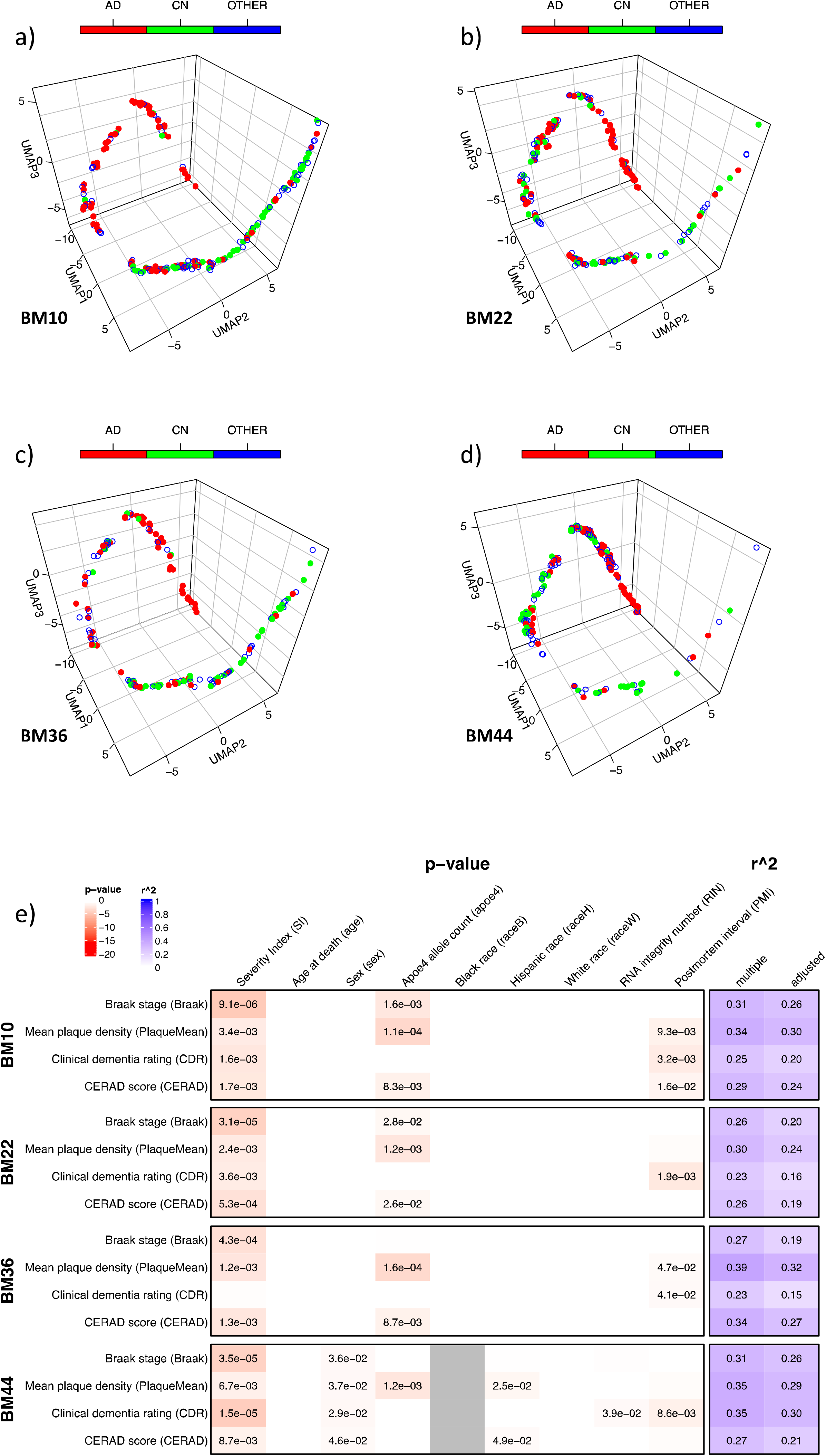
The pseudo-temporal trajectory from the transcriptomes of four brain regions of MSBB cohort and the SI correlation with phenotypic data by applying the trained deep learning model and mapping to the same 3D space as ROSMAP data. AD = Alzheimer’s disease (CDR >= 1 and Braak >= 4 and CERAD <=2); CN = control (CDR <= 0.5 and Braak <= 3 and CERAD >=3); OTHER = all other subjects. a-d) The trajectory for transcriptomes from regions BM10/20/36/44 respectively. e) The model metrics for the linear regression between SI and all the neuropathological and clinical biomarkers, by different brain regions with p values shown in cells with p < 0.05. Grey cells indicate no data. Detailed model metrics are reported in Table S6.

### Network analysis identified four major gene modules for disease progression

A clear bimodal distribution on the logarithm scale was observed in the weight distribution for the 15,582 genes in the deep learning model (Figure S2a). The cutoff was set at 1.6e-4, which generated 593 genes as the significant genes (index genes) associated with AD progression (Table S7). The distribution of these genes showed some, though not complete overlap with those differentially expressed genes (DEGs) identified in previous work for ROSMAP cohort alone (syn8456629, Figure S2b), or from the AMP-AD meta-analysis (syn11914606, Figure S2c), as unlike DEGs, some of these index genes may have smaller fold change (log2FC), or not pass the significant p value cutoff in comparison between AD vs control. Based on network analysis for these 593 genes’ expression profile, six co-expression modules were identified, with four of them showing significant correlation with multiple phenotypes (Figure 5a). Among them, the green module (n = 41) is significantly correlated with all the neuropathological and clinical biomarkers, while the turquoise module (n = 308) was found to be especially significantly correlated with tangles, and the brown module (n = 61) with amyloid. Notably, the turquoise module’s directions of correlations with the pathological traits were reversed with the other three, although all showed significant contributions to cognitive functions. The yellow module (n = 53) was only significantly correlated with the diagnosis of Parkinson’s disease and amyloid, while the grey module (n = 63) showed little correlation with any of the pathological phenotypes as expected. We also decomposed the contributions of each module to the SI and found that the SIs derived for turquoise and green modules showed strongest concordance with all the biomarkers and cognitive function score (Figure S3, Table S8).

**Figure 5.**
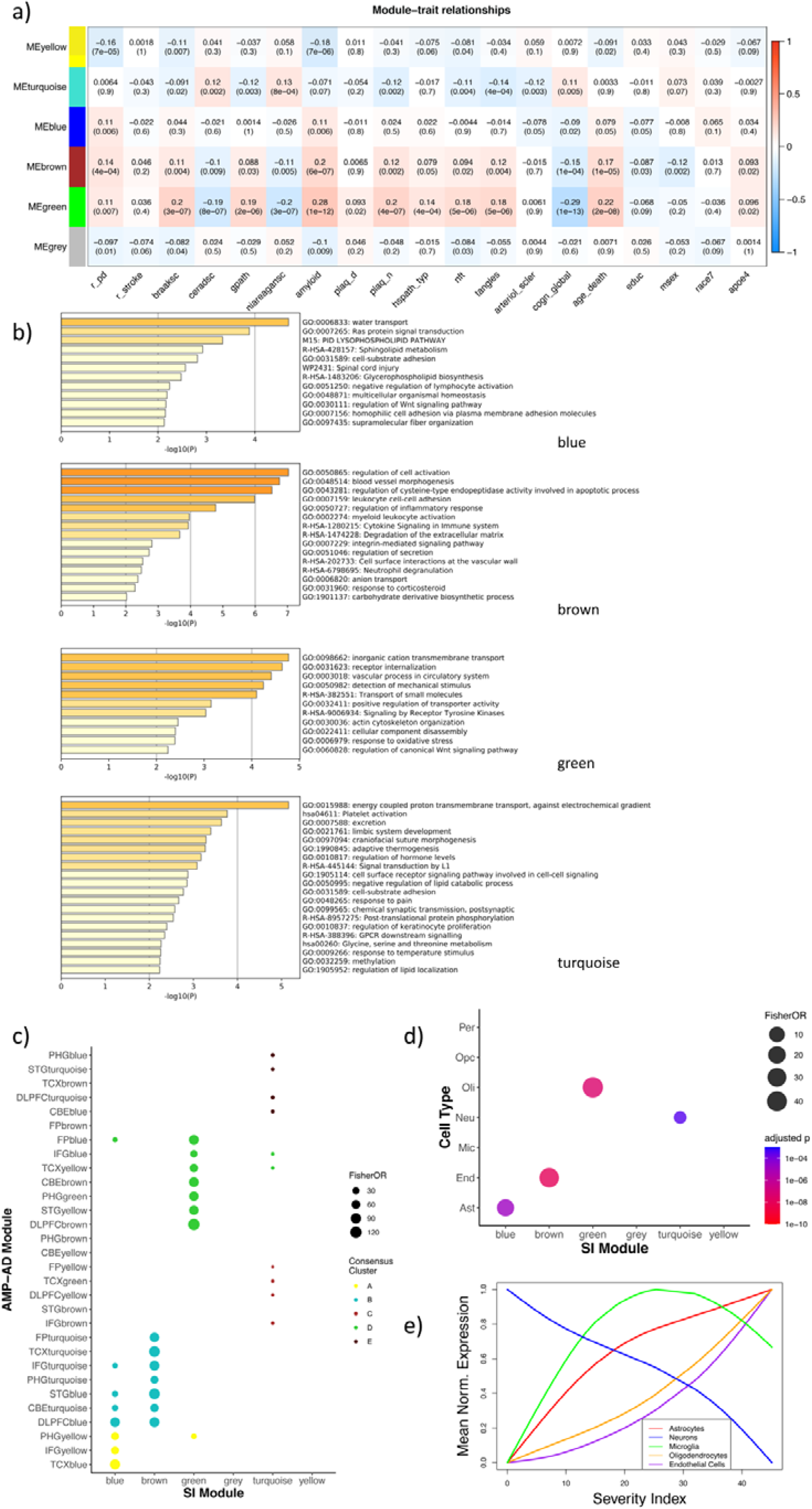
Network and gene set analysis of the index genes. a) Module trait relationship between the eigengenes of the six coexpression modules and individual traits. b) Functional enrichments for the index genes present in the four modules. c) The modules were examined for overlap (Fisher’s exact test) with the gene sets from the mega analysis of the AMP-AD consensus RNA-seq coexpression modules^23^. Overlaps were shown for those adjusted p (Bonferroni correction) < 0.001. d) Cell type enrichments using Fisher’s exact test of gene set overlap of the modules with cell type specific gene sets from human reference single-cell RNA-seq data^37, 38^ . Overlaps were shown for those adjusted p (Bonferroni correction) < 0.001. Ast = astrocytes; End = endothelial cells; Mic = microglia; Neu = neurons; Oli = oligodendrocytes; Opc = oligodendrocyte progenitor cells; Per = pericytes. e) Cell type marker gene expression signatures as a function of SI. Mean expression of cell markers for astrocytes, neurons, microglia, oligodendrocytes, and endothelial cells were plotted and colored respectively.

Functional enrichment of the genes present in the four key modules showed that they were implicated in different processes. For the turquoise module, the enriched terms were related to metabolism and hormone activities; for the brown module, they were related to vascular dysfunctions (Figure 5b). Green and blue modules were implicated in metabolic abnormalities. Interestingly, the four modules were found to overlap mostly with the consensus cluster A (for blue), cluster B (for brown), cluster D (for green) and cluster C and E (for turquoise) respectively from the meta-analysis of the AD human brain transcriptome^23^ (Figure 5c) and concordantly, the three upregulated modules were each enriched for astrocytes (blue), oligodendrocytes (green), and endothelial cells (brown), and the downregulated turquoise module was enriched in neurons (Figure 5d). Their overlaps with the curated AD gene sets from known databases and the gene sets from individual cohort study in the AMP-AD consortium confirmed this observation (Figure S5).

The change of cell type markers as a function of SI along disease progression (Figure 5e) recapitulates known cellular changes such as neurodegeneration and gliosis. In addition, it suggests that microgliosis and astrogliosis happen earlier in the disease, while oligodendrocytes and endothelial cells are activated at a faster pace at the late stage. Interestingly, marker genes in microglia don’t express monotonically in the process, which is especially prominent in males (Figure S7).

## Discussions

It is increasingly accepted that AD and Alzheimer’s Disease Related Dementias (AD/ADRD) are a spectrum of related diseases that have similar clinical and neuropathological manifestations.^39^ The progressive nature and relatively long deteriorating course warrant the study of the disease as a continuum instead of discrete states. However, the lack of longitudinal data from brain tissues of the same individuals prompts recent studies to model the gene expression dynamics of AD as pseudo-temporal trajectories, using data collected from postmortem brain tissues in large cohorts, with various degrees of neurodegeneration and dementia severity. These unsupervised machine learning approaches provide novel insights into the progressive nature of AD and demonstrate that population level cross-sectional transcriptomic data could be capitalized to capture the evolution of multiple AD-related neuropathology or cognitive impairment. In this work, we present a deep-learning framework, which includes supervised classification and unsupervised dimension reduction of transcriptomic data to derive a trajectory that strongly mirrors AD specific severity, since we have used the neuropathologically confirmed AD and control as the two termini to train the model. The SI defined by the distance along a trajectory could be utilized as a metric to evaluate the progression and staging of AD. The transcriptomic signature identified by the model sheds new light on the evolution of the gene expression profile across the disease course and illustrates the utility of deep learning approaches for the investigation of neurodegenerative diseases such as AD.

Notably, the deep learning component in our framework consists of a neural network of only three layers. There are two reasons we chose such a network. Firstly, the learning datasets from ROSMAP composed of only 243 neuropathologically confirmed AD and control subjects. Although this is one of the largest postmortem brain transcriptomic datasets of LOAD cohorts publicly available to date, more learning layers on this scale of data don’t necessarily produce better results, as 4 or more layers in the current model didn’t improve the model’s performance by our tests. Conversely, a single hidden layer in the model enables a straightforward interpretation of the embedding of the input gene features, which facilities the identification of index genes for downstream analysis. With the rapidly accelerating generation of multi-omic data for AD, this current framework could be easily extended to include additional layers for exploiting larger datasets, with both more samples or genomic features, and other phenotypic data types.

The original trajectory derived for the DLPFC tissues from the ROSMAP cohort suggests that the course of AD may be characterized by the multidimensional, nonlinear nature of transcriptome dynamics in the neurodegenerative process. The neuropathologically confirmed AD and control subjects were mostly clustered in two corners, with relatively few subjects located between the two clusters along the trajectory. When other samples with various neuropathology and/or clinical diagnosis were mapped to the same space, they were distributed widely along the trajectory, with a considerable portion in-between the two clusters. SI still significantly predicted cognitive function (p = 3.5e-7) and neuropathology (p < 7e-2) for the other samples after excluding the AD/control termini used in the training process, demonstrating the validity of the model for the general dementia population. This was further demonstrated by the application of the model to the external datasets in the AMP-AD consortium, the MAYO and MSBB RNA-seq profiles. For all the brain regions known to be susceptible to AD related pathology (TCX, FP/PHG/IFG/STG), the trajectories showed differentiating clusters of AD and control subjects. In stark contrast, for the transcriptome in cerebellum (CER), which possesses a distinct cellular architecture and is comparatively spared by AD neuropathology, the subjects’ locations are randomly distributed along the trajectory. Quantitively, SIs derived from the trajectories confirmed the observation, with all of them correlating with the biomarkers closely except those for CER. Notably, thanks to the detailed neuropathological and clinical characterizations in the ROSMAP cohort enabling a covariate analysis in the linear regression model of SI against global cognitive function (Figure 2c, Table S3), cognitive impairment in the general dementia population is also partially attributable to multiple other comorbidities such as Parkinson’s and neocortical Lewy body disease, as well as hippocampal sclerosis and arteriolosclerosis. This underscores the heterogeneity of the broad dementia spectrum and the urgent need to dissect the spectrum by distinguishing the AD specific pathology and those caused by other related diseases for precision dementia diagnosis and treatment.

Deep learning methods have been demonstrated recently to be able to capture complex, non-linear transcriptomic features that are not learned using conventional gene expression data analysis methods in AD cohorts.^40^ In our model, it searches the data for correlated features and combines them by amplifying the underlying signals with adjustable weights and the sigmoid function, so it can extract the genomic features most pertinent to the questions we are asking, i.e. the coordinated transcriptomic signature differentiating definitive AD and control. The model is completely portable and applicable to any gene expression data, and the reproducible significant results in external datasets from pathologically affected tissues manifest that the index genes indeed play significant roles in the progression of LOAD.

The index genes identified by the deep learning model have unique implications, in our understanding of AD etiology, as well as pursuit of novel therapeutics. They have some overlap, but considerably differ from those DEGs identified by differential expression (DE) analysis. Recently it has been demonstrated that certain gene expression profiles from RNA-seq experiments are generic, with a high probability of DE across a wide variety of biological conditions, so their specificity related to disease mechanisms have been challenged.^41^ Additionally, DEGs are those passing certain statistical cutoffs systematically for both fold change and p values selected as the “hit list” for further interpretation and validation. The cutoffs are arbitrarily set following a convention (e.g. fdr < 0.05, |log2FC| > 0.263 in the DEG analysis for ROSMAP dataset (syn8456629)), so it might not be able to capture subtle, intrinsic, and coordinated gene expression signatures due to disease pathology in the high dimensional data, especially from bulk tissues.^40, 42^ This is further demonstrated by the fact that neither DEGs identified by ROSMAP dataset alone (syn8456629) nor those from the AMP-AD meta-analysis (syn11914606) could fully reproduce the progressive trajectory in any of the three transcriptomic datasets, especially the external datasets when applied with the model (results not shown).

Among the six co-expression modules from the index genes, blue, brown, green, and turquoise modules are significantly correlated with AD phenotypical hallmarks, with the first three upregulated in the progression, while turquoise genes downregulated. Together with the cell type enrichment analysis showing the three modules are enriched in astrocytes, oligodendrocytes, and endothelial cells respectively, and turquoise module enriched in neurons, this is consistent with the results obtained by the recent work of neurodegeneration pseudotime estimation^5^ and cellular composition deconvolution,^42^ which shows a reduction in the neuronal populations as AD progresses, and an increase in expression associated with activation of endothelial and glial cells, as also demonstrated by the change of mean expression for the marker genes of each cell type along AD progression. Interestingly, the transcriptomic signatures obtained from our study show high similarity with the signatures obtained from a recent single-nucleus transcriptome analysis from the prefrontal cortical samples of AD patients and normal control subjects,^38^ where higher proportion of endothelial nuclei were sampled and dysregulated pathways are associated with blood vessel morphogenesis, angiogenesis and antigen presentation. Notably, these functions are also implicated in the top common blood-brain functional pathways relevant for LOAD progression in the recent study of gene expression trajectories in AD.^5^ In addition, the enriched functions of proton transport implicated in mitochondrial functions and cell signaling pathway in the turquoise module were also found to be associated with the overlapped DEGs in neurons from two independent single nuclei transcriptomic studies from AD patients.^38, 43^ Strikingly, although the brown module is mostly overlapped with the consensus module cluster B from the AMP-AD transcriptome meta-analysis^23^ (Figure 5c), it is not enriched in microglia but endothelial cells (Figure 5d). It is most likely a submodule in the cluster which represents the signatures from endothelial cells. Likewise, the turquoise module as a subset of the consensus module cluster E, shows comparatively poor enrichment for cell-type expression signatures and were not well annotated or represented among curated AD pathways (Figure S5). These results collectively suggest that the transcriptomic signatures identified by the deep learning framework constitute intrinsic molecular changes at cellular level associated with AD’s progression. Whether the changes are the drivers of the progression or just physiological responses accompanying the progression awaits further examination. With the identification of the four modules each enriched in a specific cell type and strongly associated with AD severity, the future work will be a scrutiny of these genes for their roles in AD’s progression.

It is more and more evident that sexual dimorphism plays an important role in AD’s development and progression.^44^ In the recent work by Mukherjee et al.^5^ where unsupervised learning methods were applied to the ROSMAP transcriptome data, predictive pseudotimes were only observed for female samples, highlighting great diversity of the gene expression profiles between the sexes. While we didn’t explicitly include sex as a feature node in the deep learning model, our analysis showed that sex effect has been modeled through the expression levels of sex markers such as XIST.^45^ When we compare SI with the pseudotime obtained by unsupervised learning on the same subjects as reported in the work of Mukherjee et al., higher degree of concordance is obtained for the female samples than the male (Figure S6a). SI were found to be significantly associated with the neuropathological and clinical biomarkers in both sexes, by both linear and logistic regressions (Figure S6b, c). In the plots stratified by sex of mean marker gene expression as a function of SI, we observe not only consistent overall patterns in both sexes, but also distinct curves for some cell types, e.g. neuron and microglia (Figure S7b, c). Neuronal degeneration occurs earlier and faster in female, although eventually both sexes converge at a similar total loss. For microglia, the change is not monotonic, especially in male. These observations highlight important cell type specific contributions to AD progression in different sexes, which has drawn considerable research efforts in recent years.^46, 47^ Lastly, sex has also been considered as a covariate in the linear regression of SI against all the neuropathological and clinical biomarkers in all three cohorts. They are all not significant except in the IFG region from the MSBB cohort (Figure 4e, Table S6), where female sex is associated with higher clinical and pathological severity. All of the above work collectively demonstrates that our model captures generalized transcriptomic features present in both sexes for most AD affected brain regions like DLPFC, while in specific brain region like IFG, there are additional sex effects that await further investigation.

From the trajectories derived for different AD affected brain regions and their model metrics (e.g. MSBB cohort, Figure 4, Table S6), it is evident that there are perceptible although subtle variations within their expression profiles, consistent with the observations from multiple analyses of transcriptomic data for the cohort.^48, 49^ This highlights the necessity of analyzing tissue and region-specific datasets to better understand interactions between brain region and molecular disease states within AD. For instance, with preliminary validation in this work, a similar deep learning framework could be designed specifically to dissect transcriptomes from peripheral tissues such as blood of AD cohorts to seek highly sensitive and specific targets, to complement any biomarkers currently being actively pursued for early diagnosis of the devastating disease. Similarly, the application of such an approach has broad utility for use with any high dimensional multi-omics data such as proteomics, metabolomics and epigenomics, thus opening another channel for the application of artificial intelligence in the genomics field for pursuing early diagnosis and effective treatment of neurodegenerative diseases.

## Supporting information

All supplemental tables

## Acknowledgements

The results published here are in whole or in part based on data obtained from the AD Knowledge Portal (https://adknowledgeportal.org) (syn2580853). The RNAseq Harmonization Study (rnaSeqReprocessing) was supported by the NIA grants U01AG046152, U01AG046170, U01AG046139, and U24AG061340. Study data in ROSMAP cohort were provided by the Rush Alzheimer’s Disease Center, Rush University Medical Center, Chicago. Data collection was supported through funding by NIA grants P30AG10161 (ROS), R01AG15819 (ROSMAP; genomics and RNAseq), R01AG17917 (MAP), R01AG36836 (RNAseq), the Illinois Department of Public Health and the Translational Genomics Research Institute. Additional phenotypic data were requested at https://www.radc.rush.edu. The data for MSBB cohort were generated from postmortem brain tissue collected through the Mount Sinai VA Medical Center Brain Bank and were provided by Dr. Eric Schadt from Mount Sinai School of Medicine. The MSBB study was led by Dr. Nilufer Ertekin-Taner and Dr. Steven G. Younkin, Mayo Clinic, Jacksonville, FL using samples from the Mayo Clinic Study of Aging, the Mayo Clinic Alzheimer’s Disease Research Center, and the Mayo Clinic Brain Bank. Data collection was supported through funding by NIA grants P50AG016574, R01AG032990, U01AG046139, R01AG018023, U01AG006576, U01AG006786, R01AG025711, R01AG017216, R01AG003949, NINDS grant R01NS080820, CurePSP Foundation, and support from Mayo Foundation. Study data includes samples collected through the Sun Health Research Institute Brain and Body Donation Program of Sun City, Arizona. The Brain and Body Donation Program is supported by the NINDS (U24NS072026 National Brain and Tissue Resource for Parkinson’s Disease and Related Disorders), the NIA (P30AG19610 Arizona Alzheimer’s Disease Core Center), the Arizona Department of Health Services (contract 211002, Arizona Alzheimer’s Research Center), the Arizona Biomedical Research Commission (contracts 4001, 0011, 05-901 and 1001 to the Arizona Parkinson’s Disease Consortium) and the Michael J. Fox Foundation for Parkinson’s Research. We thank the authors of DeepType for technical assistance of training the deep learning model and integrating the method into our framework. We thank the staff at Research Resource Sharing Hub, Rush Alzheimer’s Disease Center, Rush University Medical Center, Chicago for their efforts to share the unreleased data from the ROSMAP study. The authors acknowledge Research Computing at Arizona State University for providing computing resources that have contributed to the research results reported within this paper. The authors also thank the constructive suggestions from the reviewers for improvement of this manuscript.

## Supplementary material

Supplementary material is available at Brain online.

---

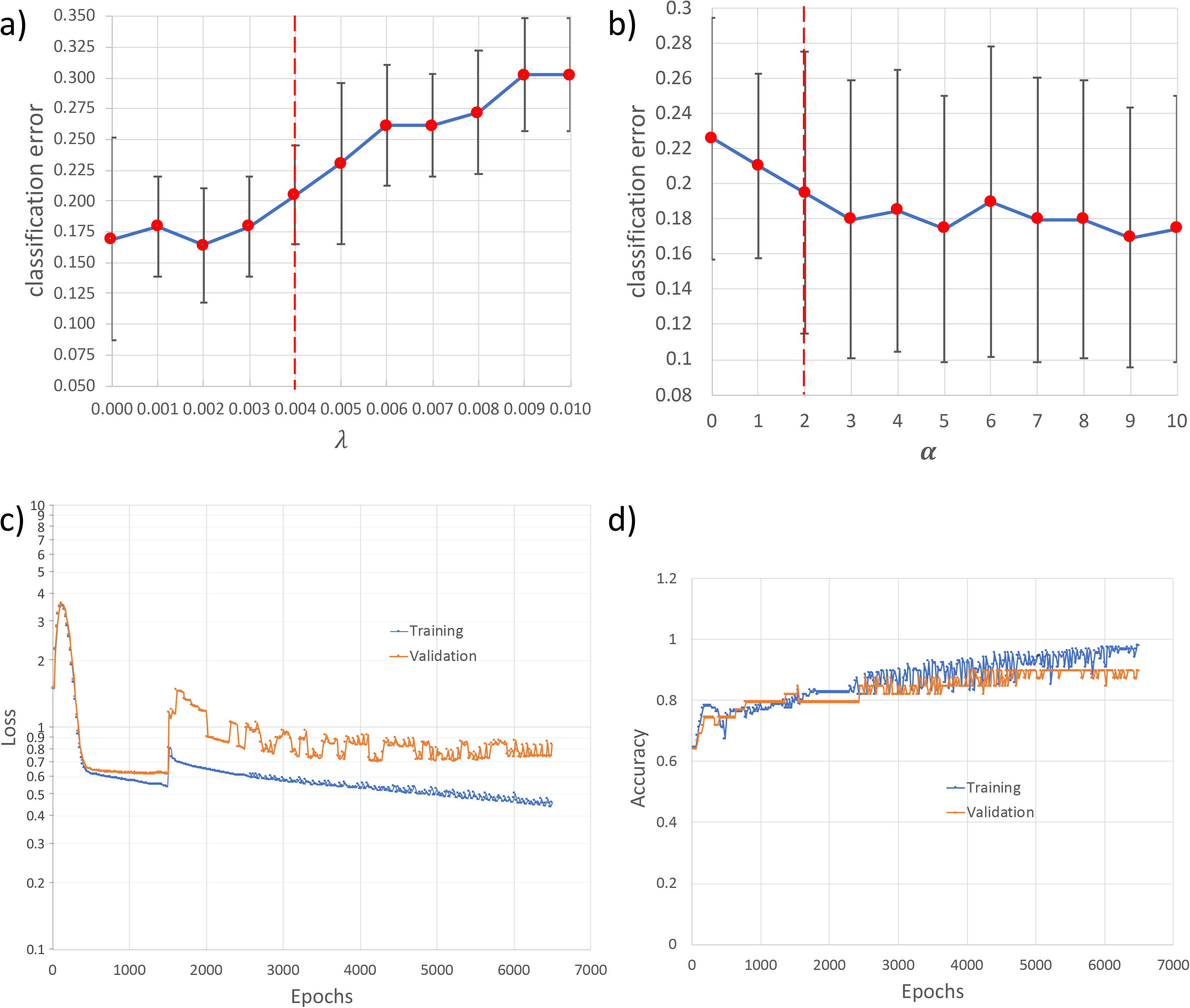

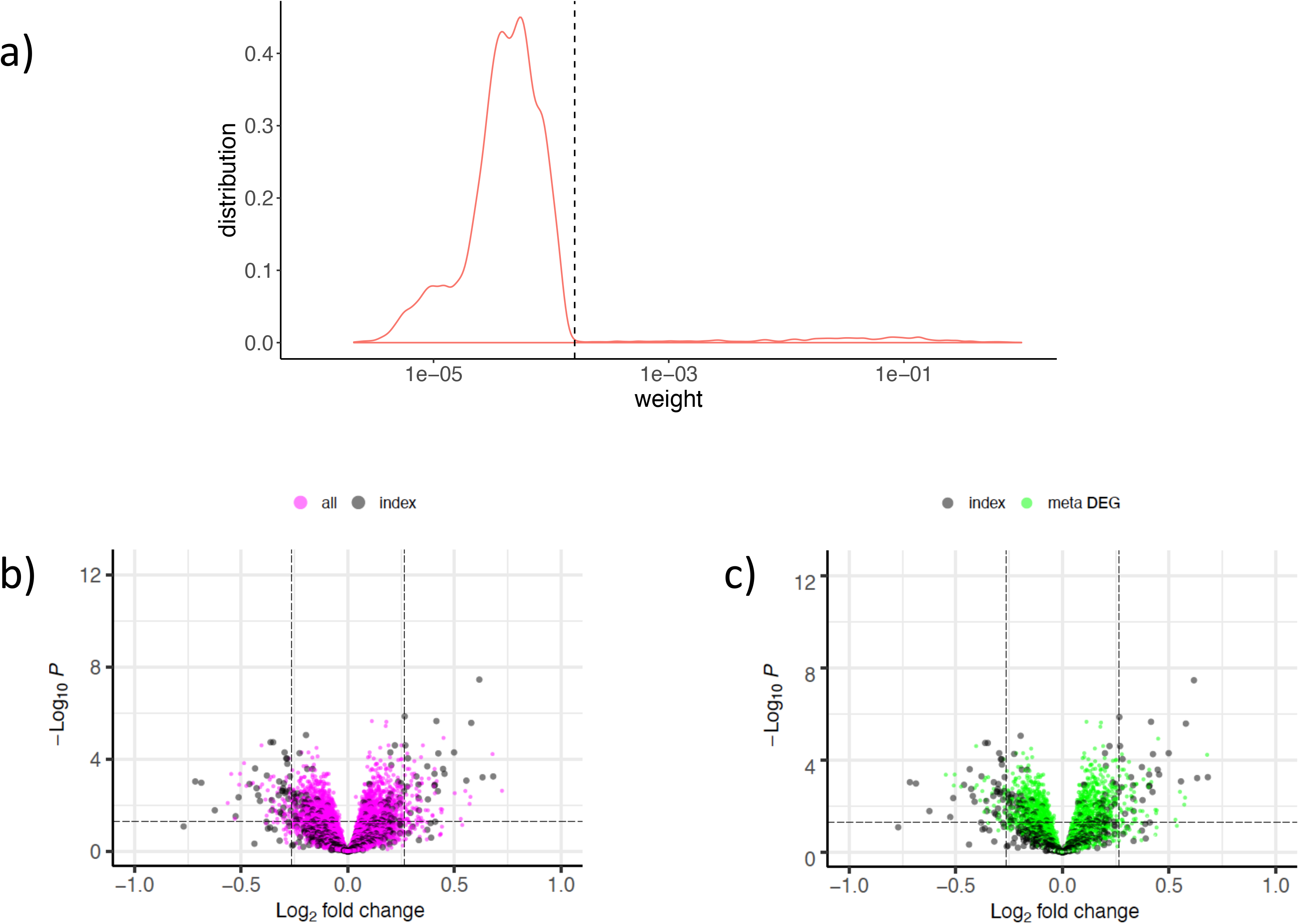

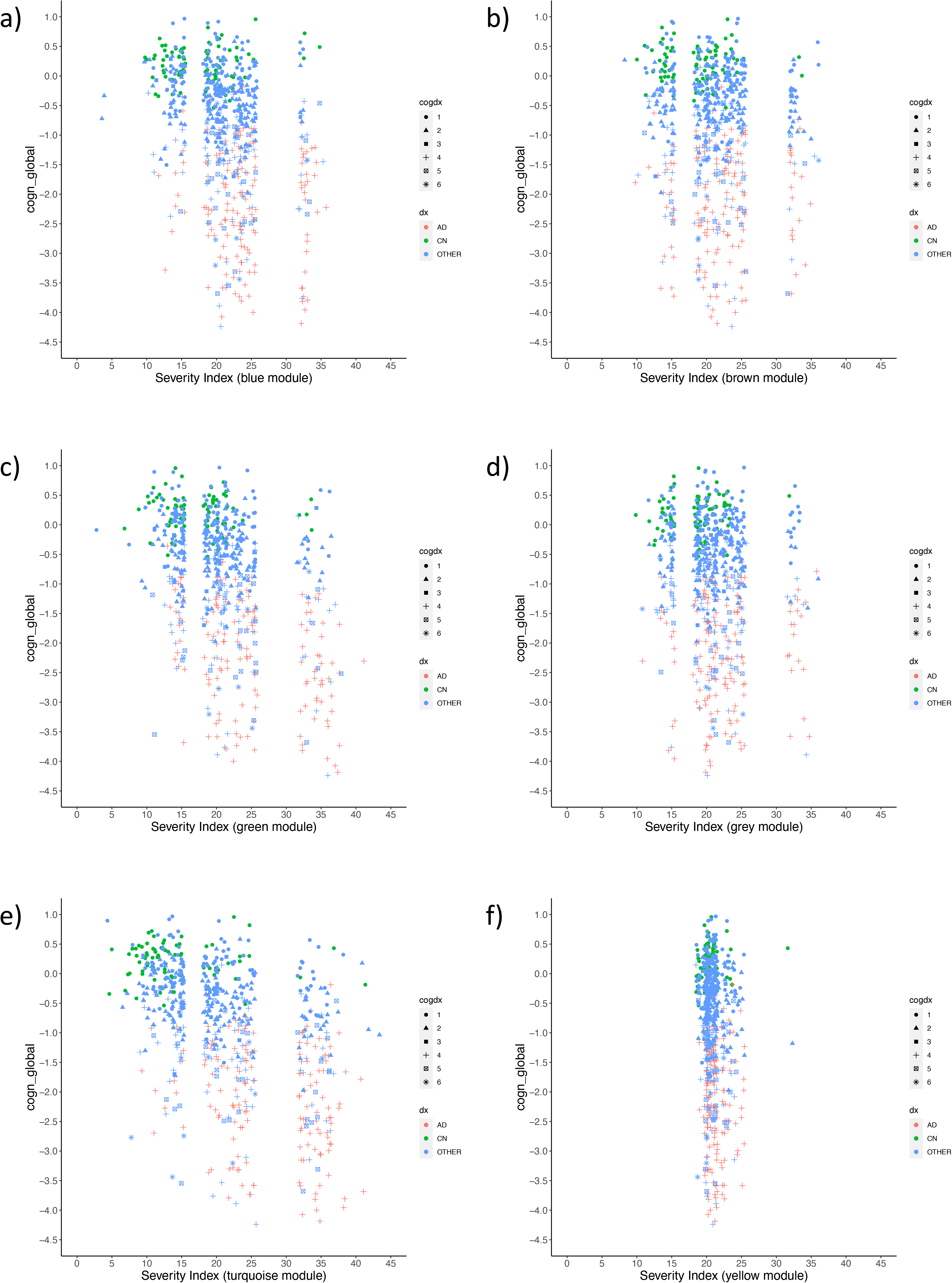

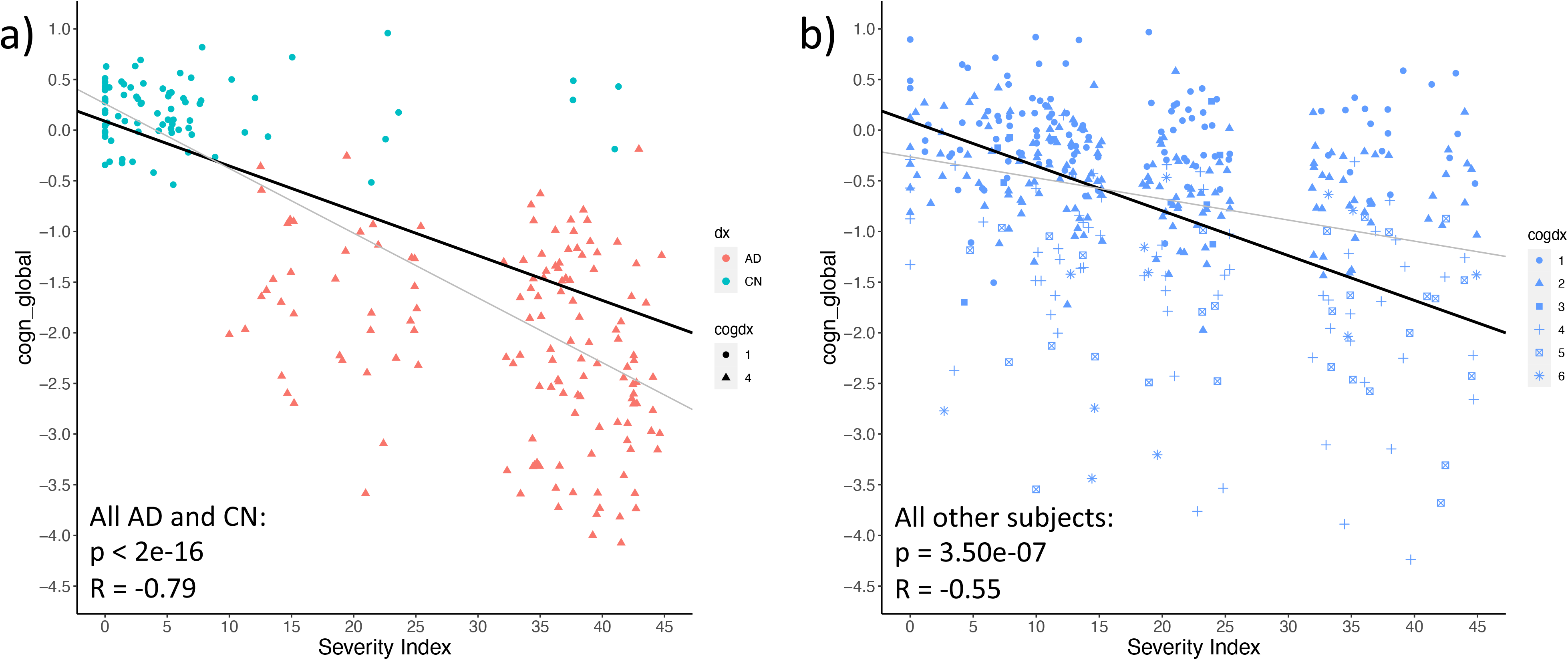

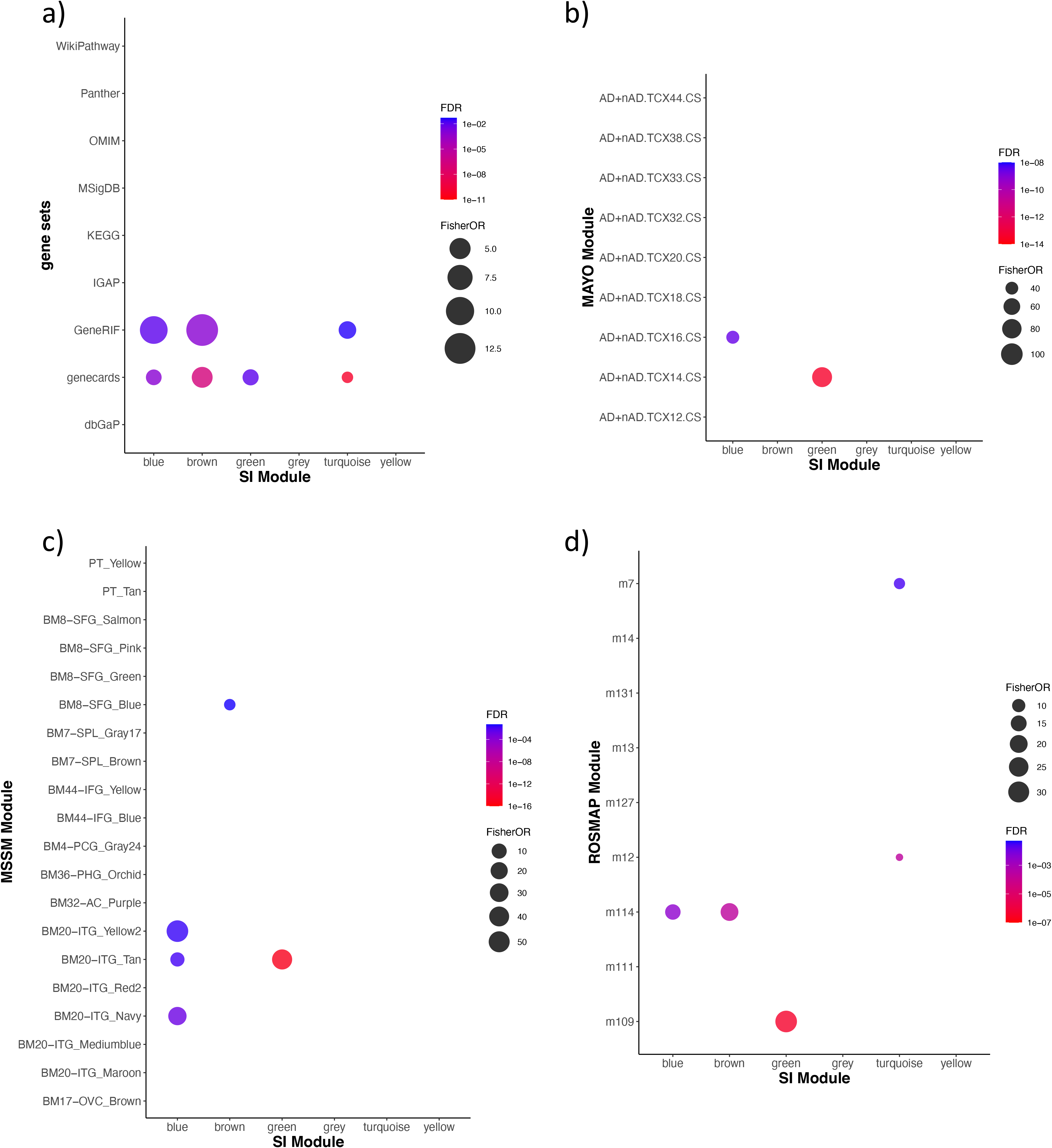

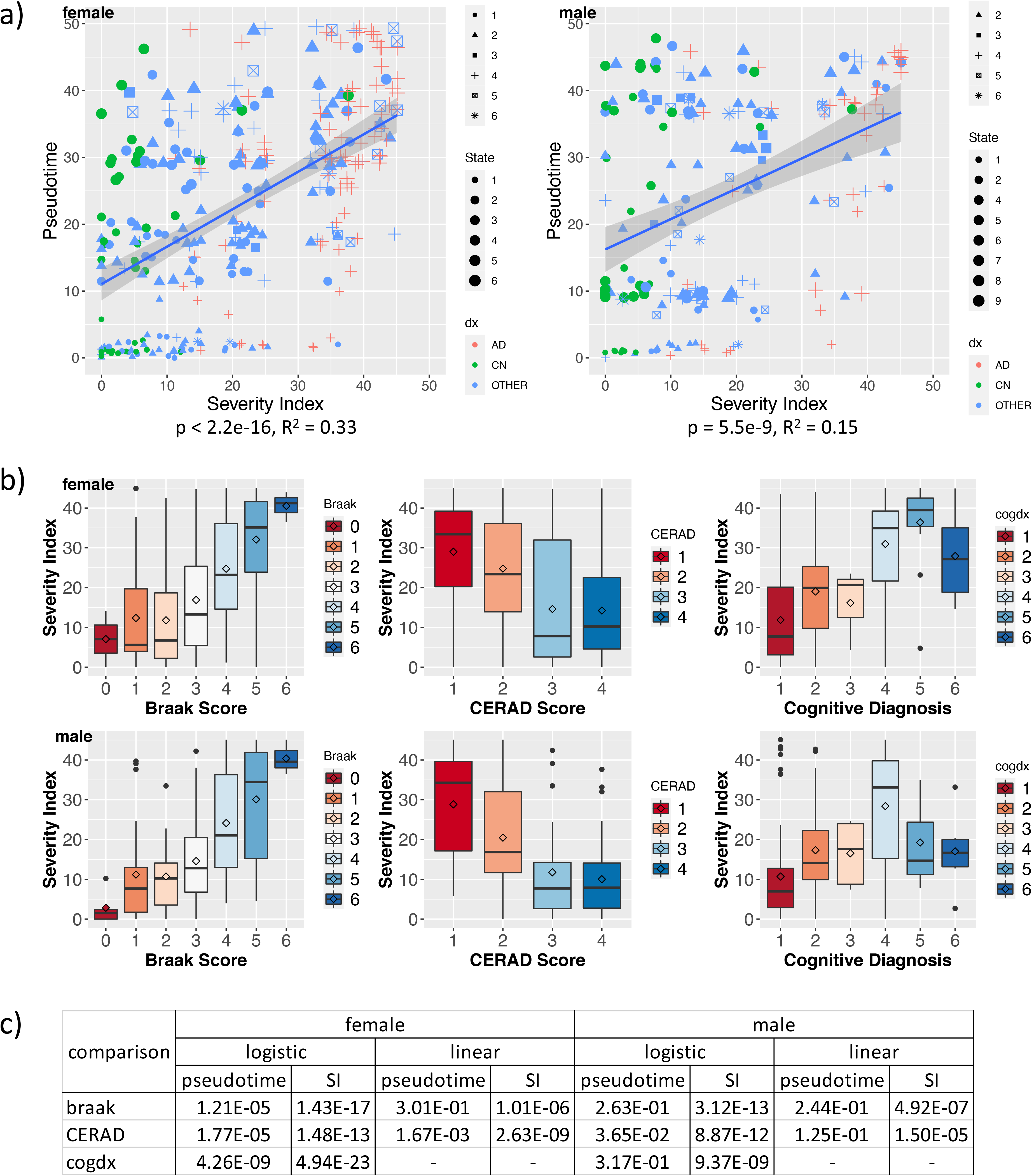

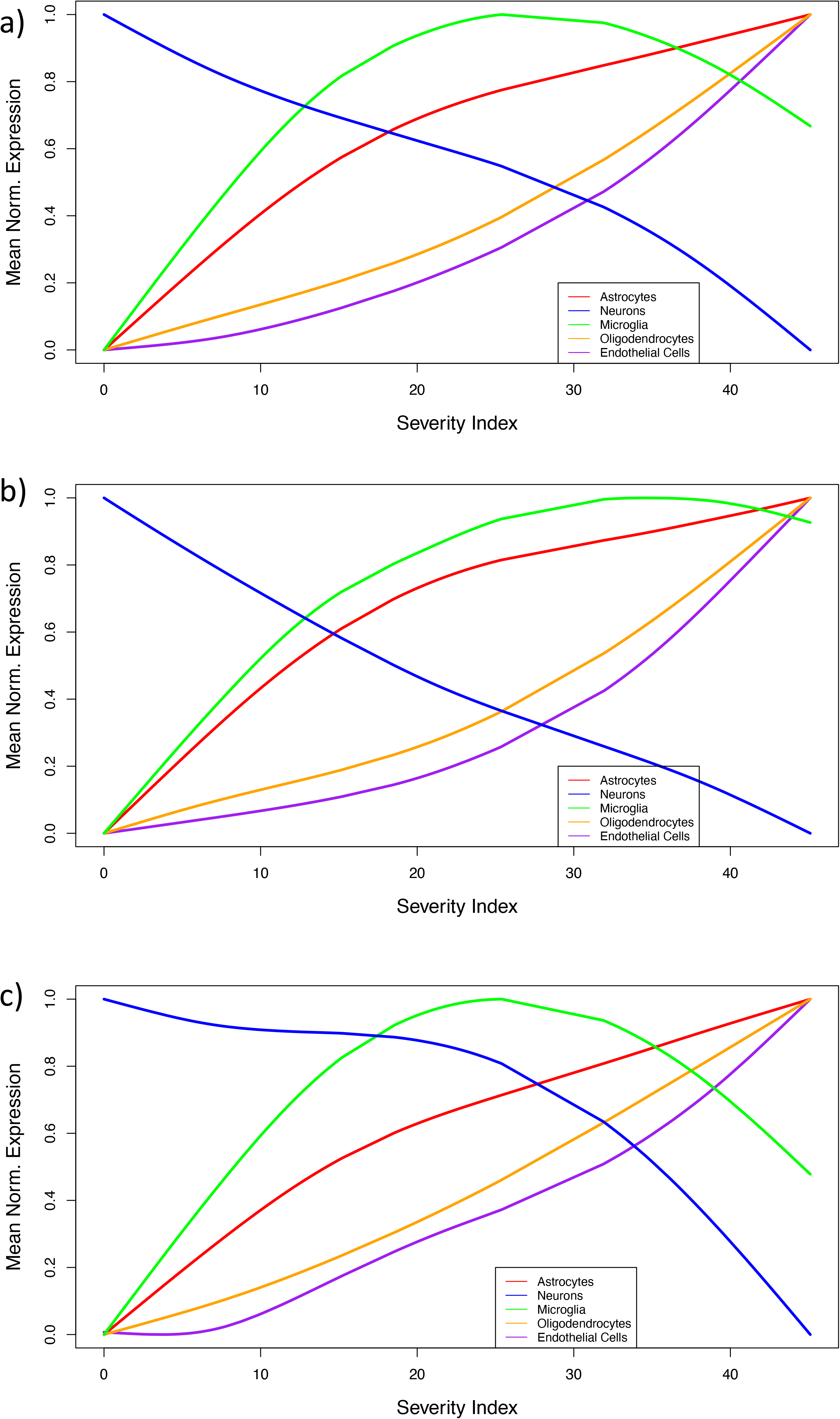

