## Supplementary material for "Deep learning-based brain transcriptomic signatures associated with the neuropathological and clinical severity of Alzheimer’s disease": All supplemental tables

**Table S1 Demographic information for all the subjects included in RNA-seq data used in this study.**

| Study | Sample size# | Tissue | AD | | control | | other** | |
|---|---|---|---|---|---|---|---|---|
| | | | gender (M/F) | age* (mean/SD) | gender (M/F) | age* (mean/SD) | gender (M/F) | age* (mean/SD) |
| ROSMAP | 634 | DLPFC | 46/110 | 91.0/5.7 | 39/48 | 84.1/6.8 | 143/248 | 88.8/6.5 |
| MAYO | 263 | CER | 32/47 | 82.5/7.7 | 38/36 | 82.4/8.3 | 61/49 | 76.7/7.6 |
| | 266 | TCX | 31/49 | 82.6/7.7 | 38/36 | 82.7/8.4 | 62/50 | 77.0/7.8 |
| MSBB | 214 | FP (BM10) | 27/58 | 84.4/6.6 | 25/29 | 80.6/8.6 | 21/54 | 85.5/6.0 |
| | 191 | STG (BM22) | 29/50 | 83.0/7.2 | 19/26 | 80.1/9.4 | 17/50 | 85.6/6.2 |
| | 162 | PHG (BM36) | 18/44 | 84.4/6.9 | 23/21 | 79.5/9.5 | 17/39 | 84.6/6.7 |
| | 187 | IFG (BM44) | 24/49 | 84.0/6.9 | 23/21 | 80.1/9.2 | 18/52 | 85.6/6.1 |

* All those with age > 90 in MAYO/MSBB cohort were calculated as 90

** All diagnosis confirmed by neuropathological assessments

### Sample size after outlier removal in RNA-seq normalization

**Table S2 Description of the variables used in the linear regression model for ROSMAP cohort.**

| category | | code | description | value range | note |
|---|---|---|---|---|---|
| diagnosis | | cogdx | Final consensus cognitive diagnosis | 1,2,3,4,5,6 | Clinical consensus diagnosis of cognitive status at time of death |
| variables used in linear regression model | dependent variable, AD-related quantitative measurements | braaksc | Braak stage | 0,1,2,3,4,5,6 | Semiquantitative measure of neurofibrillary tangles |
| | | ceradsc | CERAD score | 1,2,3,4 | Semiquantitative measure of neuritic plaques |
| | | niareagansc | NIA-Reagan diagnosis of AD | 1,2,3,4 | Consensus postmortem diagnosis based on both neurofibrillary tangles (Braak) and neuritic plaques (CERAD) |
| | | gpath | Global AD pathology burden | 0.000 ~ 3.200 | Quantitative summary of three AD pathologies: neuritic plaques (n), diffuse plaques (d), and neurofibrillary tangles (nft) |
| | | amyloid | Overall amyloid level | 0.000 ~ 19.925 | Overall amyloid level - Mean of 8 brain regions |
| | | plaq_d | Diffuse plaque burden | 0.000 ~ 4.912 | Diffuse plaque summary based on 5 regions |
| | | plaq_n | Neuritic plaque burden | 0.000 ~ 5.004 | Neuritic plaque summary based on 5 regions |
| | | nft | Neurofibrillary tangle burden | 0.000 ~ 6.099 | Neurofibrillary tangle summary based on 5 regions |
| | | tangles | Tangle density | 0.000 ~ 78.523 | Tangle density - Mean of 8 brain regions |
| | | cogn_global | Global cognitive function | -4.239 ~ 1.224 | Z score from 19 cognitive tests |
| | independent variable, non-AD quantitative measurements | SI | Severity Index | 0.00 ~ 45.10 | defined in this work |
| | | age_death | Age at death | 67.37 ~ 108.28 | |
| | | educ | Year of education | 3 ~ 28 | |
| | | msex | Sex | 0,1 | |
| | | race7 | Racial group | 1,2,3,4,5,6,7 | categorical |
| | | apoe4 | Apoe4 allele count | 0,1,2 | |
| | | RIN | RNA integrity number | 5.0 ~ 9.9 | |
| | | PMI | Postmortem interval | 1.0 ~ 40.8 | |
| | | r_pd | Clinical diagnosis of Parkinson's Disease | 1,2,3,4 | |
| | | r_stroke | Clinical stroke diagnosis | 1,2,3,4 | |
| | | dlbdx | Lewy body disease | 0,1,2,3 | categorical; 4 stages of pathologic diagnosis of Lewy body diseases |
| | | hspath_typ | Hippocampal sclerosis | 0,1 | Definite presence of typical hippocampal sclerosis |
| | | arteriol_scler | Arteriolosclerosis | 0,1,2,3 | 4 stages of arteriolosclerosis |

**Table S3 Model metrics for the linear regression between global cognitive function and the dependent variables (as defined in Table S2) stratified by diagnosis groups in ROSMAP DLPFC samples. Cells with significant p values (< 0.05) are shown in bold.**

| target = global cognitive function | all subjects (n = 634) | | | | | | all OTHER subjects (n = 391) | | | | | | all AD and CN (n = 243) | | | | | |
|---|---|---|---|---|---|---|---|---|---|---|---|---|---|---|---|---|---|---|
| | Estimate | Std. Error | t value | Pr(>\|t\|) | PVE | sig. code | Estimate | Std. Error | t value | Pr(>\|t\|) | PVE | sig. code | Estimate | Std. Error | t value | Pr(>\|t\|) | PVE | sig. code |
| (Intercept) | -0.347 | 0.779 | -0.445 | 6.56E-01 | | | -1.997 | 0.937 | -2.131 | **3.41E-02** | | * | 1.683 | 1.349 | 1.247 | 2.14E-01 | | |
| SI | -0.036 | 0.003 | -11.345 | **< 2E-16** | 34.96 | *** | -0.021 | 0.004 | -5.237 | **3.50E-07** | 15.23 | *** | -0.050 | 0.005 | -9.540 | **< 2E-16** | 57.45 | *** |
| age_death | -0.011 | 0.007 | -1.654 | 9.89E-02 | 0.47 | . | -0.002 | 0.008 | -0.266 | 7.91E-01 | 0.17 | | -0.028 | 0.012 | -2.374 | **1.89E-02** | 1.60 | * |
| educ | 0.034 | 0.012 | 2.787 | **5.57E-03** | 0.20 | ** | 0.024 | 0.015 | 1.645 | 1.01E-01 | 0.07 | | 0.056 | 0.022 | 2.575 | **1.11E-02** | 1.43 | * |
| msex | -0.099 | 0.082 | -1.209 | 2.28E-01 | 0.38 | | -0.059 | 0.097 | -0.610 | 5.43E-01 | 0.37 | | -0.050 | 0.141 | -0.353 | 7.25E-01 | 0.05 | |
| race7 | -0.018 | 0.303 | -0.060 | 9.52E-01 | 0.08 | | -0.278 | 0.330 | -0.841 | 4.01E-01 | 0.35 | | 0.072 | 0.602 | 0.119 | 9.06E-01 | 0.02 | |
| apoe4 | -0.387 | 0.084 | -4.627 | **5.00E-06** | 3.48 | *** | -0.262 | 0.102 | -2.575 | **1.06E-02** | 2.82 | * | -0.323 | 0.139 | -2.314 | **2.21E-02** | 1.33 | * |
| RIN | -0.041 | 0.042 | -0.984 | 3.25E-01 | 0.37 | | 0.042 | 0.049 | 0.853 | 3.95E-01 | 0.04 | | -0.108 | 0.075 | -1.442 | 1.52E-01 | 1.05 | |
| PMI | 0.007 | 0.009 | 0.835 | 4.04E-01 | 0.00 | | 0.014 | 0.011 | 1.323 | 1.87E-01 | 0.08 | | 0.004 | 0.015 | 0.256 | 7.99E-01 | 0.01 | |
| r_pd | 0.262 | 0.052 | 5.027 | **7.49E-07** | 4.21 | *** | 0.281 | 0.058 | 4.804 | **2.70E-06** | 8.17 | *** | 0.163 | 0.096 | 1.690 | 9.33E-02 | 0.77 | . |
| r_stroke | 0.090 | 0.059 | 1.539 | 1.25E-01 | 0.19 | | 0.111 | 0.062 | 1.791 | 7.46E-02 | 0.94 | . | 0.078 | 0.126 | 0.618 | 5.38E-01 | 0.01 | |
| dlbdx3 | -0.464 | 0.133 | -3.484 | **5.49E-04** | 1.60 | *** | -0.372 | 0.169 | -2.193 | **2.92E-02** | 1.43 | * | -0.515 | 0.202 | -2.552 | **1.18E-02** | 1.72 | * |
| hspath_typ | -0.555 | 0.168 | -3.307 | **1.03E-03** | 1.43 | ** | -0.873 | 0.228 | -3.827 | **1.64E-04** | 4.15 | *** | -0.180 | 0.245 | -0.738 | 4.62E-01 | 0.12 | |
| arteriol_scler | -0.054 | 0.038 | -1.406 | 1.60E-01 | 0.26 | | -0.039 | 0.046 | -0.847 | 3.98E-01 | 0.19 | | -0.038 | 0.063 | -0.597 | 5.52E-01 | 0.09 | |
| Multiple R^2 | 0.476 | | | | | | 0.340 | | | | | | 0.656 | | | | | |
| Adjusted R^2 | 0.457 | | | | | | 0.300 | | | | | | 0.620 | | | | | |

Signif. codes: 0 ‘***’ 0.001 ‘**’ 0.01 ‘*’ 0.05 ‘.’ 0.1 ‘ ’ 1

**Table S4 Model metrics (p values and correlation coefficients) for the linear regression between all the neuropathological biomarkers and the dependent variables (as defined in Table S2) stratified by diagnosis groups in ROSMAP DLPFC samples. Cells with significant p values (< 0.05) are shown in bold.**

| target | braaksc | | | ceradsc | | | niareagansc | | | gpath | | | amyloid | | |
|---|---|---|---|---|---|---|---|---|---|---|---|---|---|---|---|
| p value | all | OTHER | AD/ CN | all | OTHER | AD/ CN | all | OTHER | AD/ CN | all | OTHER | AD/ CN | all | OTHER | AD/ CN |
| (Intercept) | 1.40E-01 | 9.55E-01 | 1.56E-01 | **5.53E-10** | **6.98E-05** | **5.06E-05** | **2.46E-12** | **1.87E-07** | **7.28E-05** | 2.56E-01 | 6.72E-01 | 8.26E-01 | 4.92E-01 | 8.94E-01 | 9.73E-01 |
| SI | **1.00E-14** | 7.29E-02 | **6.33E-16** | **2.04E-14** | **4.25E-03** | **1.21E-15** | **2.00E-16** | **1.13E-02** | **2.00E-16** | **2.00E-16** | **2.25E-03** | **7.31E-16** | **7.16E-12** | **2.54E-02** | **1.32E-12** |
| age_death | **2.71E-08** | **1.39E-04** | **1.22E-05** | **1.50E-02** | 2.93E-01 | **1.73E-02** | **7.65E-03** | **2.89E-02** | 1.80E-01 | 2.17E-01 | 5.58E-01 | 3.55E-01 | **4.39E-02** | 4.45E-01 | **3.61E-02** |
| educ | 8.39E-02 | 9.18E-02 | 1.57E-01 | 2.45E-01 | 1.81E-01 | 5.31E-01 | 7.31E-01 | 3.31E-01 | 3.05E-01 | 8.36E-01 | 9.74E-01 | 6.14E-01 | **1.90E-02** | 1.43E-01 | **1.80E-02** |
| msex | 1.79E-01 | 1.63E-01 | 3.15E-01 | 5.04E-01 | 9.98E-01 | 3.65E-01 | 9.30E-01 | 6.51E-01 | 8.06E-01 | 7.30E-01 | 4.42E-01 | 9.18E-01 | 7.01E-01 | 9.37E-01 | 8.84E-01 |
| race7 | 7.31E-01 | 6.63E-01 | 4.88E-01 | 4.24E-01 | 8.64E-01 | 4.63E-01 | 5.10E-01 | 5.91E-01 | 6.48E-01 | 8.93E-01 | 5.29E-01 | 7.63E-01 | 3.03E-01 | 6.52E-01 | 6.97E-01 |
| apoe4 | **5.66E-05** | **2.82E-02** | **2.57E-02** | **2.28E-07** | **1.45E-04** | **1.59E-03** | **3.26E-07** | **3.56E-04** | **2.88E-02** | **3.71E-12** | **1.84E-06** | **1.62E-05** | **8.60E-08** | **2.69E-04** | **3.97E-04** |
| RIN | 3.20E-01 | 1.00E+00 | 2.73E-01 | 9.08E-02 | 5.49E-01 | 1.05E-01 | 3.08E-01 | 8.45E-01 | 4.47E-01 | **6.14E-02** | 4.90E-01 | 8.61E-02 | 9.92E-01 | 7.24E-01 | 8.13E-01 |
| PMI | 1.21E-01 | **5.99E-02** | 7.82E-01 | **4.96E-02** | 1.28E-01 | 3.42E-01 | **4.84E-02** | 8.32E-02 | 3.72E-01 | **3.63E-02** | **1.62E-02** | 8.77E-01 | 3.81E-01 | 1.14E-01 | 2.08E-01 |
| r_pd | 6.10E-01 | 1.48E-01 | 4.79E-01 | 8.19E-01 | **6.85E-02** | **7.76E-03** | 9.65E-01 | 4.20E-01 | 4.90E-01 | 8.99E-01 | 1.09E-01 | 9.52E-02 | 1.72E-01 | **1.50E-02** | 2.64E-01 |
| r_stroke | 3.02E-01 | 3.19E-01 | 1.96E-01 | 6.28E-01 | 3.36E-01 | 2.24E-01 | 8.26E-01 | 8.89E-01 | 3.02E-01 | 8.13E-01 | 5.48E-01 | 2.59E-01 | 3.78E-01 | 2.21E-01 | 3.63E-01 |
| dlbdx3 | 7.54E-01 | 2.11E-01 | **2.37E-02** | 6.43E-01 | 5.69E-01 | 2.82E-01 | 8.12E-01 | 9.61E-01 | 1.00E+00 | 6.58E-01 | 7.63E-01 | 9.23E-01 | 4.20E-01 | 3.37E-01 | 5.37E-01 |
| hspath_typ | 2.49E-01 | **1.04E-02** | 6.98E-01 | 3.86E-01 | 1.69E-01 | 9.74E-01 | 9.89E-01 | 8.64E-01 | 5.67E-01 | 9.97E-01 | 8.43E-01 | 8.90E-01 | 5.11E-01 | 9.37E-01 | 1.46E-01 |
| arteriol_scler | 9.66E-01 | 2.28E-01 | 7.55E-02 | 4.99E-01 | **2.03E-02** | 7.23E-02 | 3.13E-01 | **2.22E-02** | 3.66E-01 | 2.59E-01 | 5.37E-02 | 5.99E-01 | 5.35E-02 | 1.19E-01 | 1.41E-01 |
| Multiple R^2 | 0.33 | 0.19 | 0.67 | 0.27 | 0.15 | 0.64 | 0.31 | 0.14 | 0.63 | 0.32 | 0.16 | 0.61 | 0.26 | 0.15 | 0.57 |
| Adjusted R^2 | 0.30 | 0.14 | 0.63 | 0.24 | 0.09 | 0.60 | 0.28 | 0.08 | 0.59 | 0.30 | 0.11 | 0.57 | 0.23 | 0.10 | 0.53 |

| target | plaq_d | | | plaq_n | | | nft | | | tangles | | | cogn_global | | |
|---|---|---|---|---|---|---|---|---|---|---|---|---|---|---|---|
| p value | all | OTHER | AD/ CN | all | OTHER | AD/ CN | all | OTHER | AD/ CN | all | OTHER | AD/ CN | all | OTHER | AD/ CN |
| (Intercept) | 6.49E-01 | 9.46E-01 | 8.10E-01 | 2.18E-01 | 4.76E-01 | 9.58E-01 | 1.99E-01 | 4.00E-01 | 5.22E-01 | 2.19E-01 | 4.73E-01 | 2.99E-01 | 6.56E-01 | **3.41E-02** | 2.14E-01 |
| SI | **3.24E-06** | **4.61E-02** | **4.27E-05** | **2.00E-16** | **3.64E-03** | **3.78E-16** | **2.00E-16** | **1.26E-02** | **1.08E-12** | **2.00E-16** | **1.31E-02** | **9.39E-14** | **2.00E-16** | **3.50E-07** | **2.00E-16** |
| age_death | 2.49E-01 | 9.01E-01 | 1.60E-01 | 6.31E-01 | 9.97E-01 | 7.30E-01 | 1.14E-01 | **4.47E-02** | 6.15E-01 | **1.90E-03** | **6.60E-03** | **2.14E-02** | 9.89E-02 | 7.91E-01 | **1.89E-02** |
| educ | 2.53E-01 | 6.18E-01 | 4.55E-01 | 9.43E-01 | 4.45E-01 | 2.46E-01 | **2.56E-02** | 7.10E-02 | 2.35E-01 | 1.47E-01 | 9.32E-01 | **3.68E-02** | **5.57E-03** | 1.01E-01 | **1.11E-02** |
| msex | 9.74E-01 | 5.09E-01 | 4.16E-01 | 7.61E-01 | 1.83E-01 | 6.13E-01 | 3.20E-01 | 5.51E-02 | 7.94E-01 | **3.26E-02** | **7.63E-03** | 5.88E-01 | 2.28E-01 | 5.43E-01 | 7.25E-01 |
| race7 | 9.02E-01 | 3.62E-01 | 2.34E-01 | 3.70E-01 | 7.16E-01 | 9.57E-01 | 8.13E-01 | 5.14E-01 | 6.26E-01 | 4.82E-01 | 2.62E-01 | 6.46E-01 | 9.52E-01 | 4.01E-01 | 9.06E-01 |
| apoe4 | **6.30E-07** | **5.59E-04** | **4.60E-04** | **1.47E-09** | **1.61E-06** | **2.03E-03** | **4.88E-08** | **3.74E-03** | **8.14E-04** | **1.24E-04** | 1.52E-01 | **6.77E-03** | **5.00E-06** | **1.06E-02** | **2.21E-02** |
| RIN | 3.17E-01 | 6.73E-01 | 4.28E-01 | **7.75E-02** | 6.54E-01 | **6.04E-02** | 1.10E-01 | 5.56E-01 | 2.27E-01 | 1.95E-01 | 7.76E-01 | 6.85E-02 | 3.25E-01 | 3.95E-01 | 1.52E-01 |
| PMI | 1.71E-01 | 1.60E-01 | 6.49E-01 | **6.76E-03** | **2.89E-03** | 9.67E-01 | 3.22E-01 | 1.62E-01 | 5.83E-01 | 3.14E-01 | 1.38E-01 | 5.66E-01 | 4.04E-01 | 1.87E-01 | 7.99E-01 |
| r_pd | 2.83E-01 | 8.43E-01 | **3.82E-02** | 4.76E-01 | **7.77E-03** | **2.57E-02** | 2.92E-01 | 1.53E-01 | 6.75E-01 | 7.79E-01 | 4.33E-01 | 3.60E-01 | **7.49E-07** | **2.70E-06** | 9.33E-02 |
| r_stroke | 5.23E-01 | 6.00E-01 | **6.96E-03** | 3.13E-01 | 2.45E-01 | 8.35E-01 | 5.63E-01 | 8.85E-01 | 7.61E-01 | 9.99E-01 | 5.20E-01 | 9.95E-01 | 1.25E-01 | 7.46E-02 | 5.38E-01 |
| dlbdx3 | 7.33E-01 | 6.73E-01 | 7.62E-01 | 2.60E-01 | 8.34E-01 | 1.11E-01 | 7.50E-01 | 4.99E-01 | 3.53E-01 | 4.73E-01 | 9.60E-01 | 2.38E-01 | **5.49E-04** | **2.92E-02** | **1.18E-02** |
| hspath_typ | 8.19E-01 | 4.83E-01 | 3.99E-01 | 5.33E-01 | 1.23E-01 | 2.98E-01 | 6.49E-01 | 8.22E-02 | 9.13E-01 | 1.66E-01 | **2.31E-02** | 4.49E-01 | **1.03E-03** | **1.64E-04** | 4.62E-01 |
| arteriol_scler | **5.68E-02** | **1.87E-02** | 8.92E-01 | 4.92E-01 | 9.93E-02 | 4.61E-01 | 8.08E-01 | 8.31E-01 | 4.08E-01 | 9.43E-01 | 4.19E-01 | 6.78E-01 | 1.60E-01 | 3.98E-01 | 5.52E-01 |
| Multiple R^2 | 0.16 | 0.10 | 0.42 | 0.29 | 0.20 | 0.57 | 0.30 | 0.15 | 0.52 | 0.30 | 0.16 | 0.56 | 0.48 | 0.34 | 0.66 |
| Adjusted R^2 | 0.13 | 0.05 | 0.36 | 0.26 | 0.15 | 0.53 | 0.27 | 0.10 | 0.47 | 0.27 | 0.11 | 0.51 | 0.46 | 0.30 | 0.62 |

**Table S5 Model metrics (p values and correlation coefficients) for the linear regression between neuropathological biomarkers and the dependent variables for the two brain regions in Mayo samples. Cells with significant p values (< 0.05) are shown in bold.**

| brain region | TCX | | CER | |
|---|---|---|---|---|
| target | Braak | Thal | Braak | Thal |
| (Intercept) | **8.24E-07** | **2.36E-03** | **1.96E-02** | 2.04E-01 |
| SI | **4.88E-05** | **1.56E-03** | 6.47E-01 | 4.61E-01 |
| age_death | **2.27E-05** | 1.69E-01 | **2.87E-05** | **3.94E-03** |
| gender | 9.81E-01 | 8.18E-01 | 8.07E-01 | 8.60E-01 |
| apoe4 | **1.31E-04** | **1.48E-05** | **1.19E-05** | **2.45E-06** |
| RIN | **9.15E-07** | **4.75E-04** | 8.05E-02 | 5.11E-01 |
| PMI | **1.12E-02** | **3.67E-02** | **1.24E-02** | 8.52E-02 |
| Multiple R^2 | 0.49 | 0.50 | 0.31 | 0.36 |
| Adjusted R^2 | 0.46 | 0.46 | 0.26 | 0.31 |

**Table S6 Model metrics (p values and correlation coefficients) for the linear regression between neuropathological and clinical biomarkers and the dependent variables for the four brain regions in MSBB samples. Cells with significant p values (< 0.05) are shown in bold.**

| brain region | BM10 (FP) | | | | BM22 (STG) | | | | BM36 (PHG) | | | | BM44 (IFG) | | | |
|---|---|---|---|---|---|---|---|---|---|---|---|---|---|---|---|---|
| target | Braak | Plaque Mean | CDR | CERAD | Braak | Plaque Mean | CDR | CERAD | Braak | Plaque Mean | CDR | CERAD | Braak | Plaque Mean | CDR | CERAD |
| (Intercept) | 9.24E-01 | 3.91E-01 | 7.52E-01 | **2.61E-02** | 8.78E-01 | 6.30E-01 | 1.06E-01 | 5.90E-02 | 9.43E-01 | 7.98E-01 | 3.79E-01 | **3.35E-02** | 1.65E-01 | 1.85E-01 | **4.55E-02** | **4.75E-02** |
| SI | **9.11E-06** | **3.42E-03** | **1.61E-03** | **1.72E-03** | **3.09E-05** | **2.36E-03** | **3.60E-03** | **5.28E-04** | **4.29E-04** | **1.17E-03** | 6.74E-02 | **1.28E-03** | **3.51E-05** | **6.69E-03** | **1.52E-05** | **8.74E-03** |
| age | 2.08E-01 | 9.61E-01 | 9.40E-01 | 3.68E-01 | 6.58E-01 | 8.36E-01 | 2.51E-01 | 8.80E-01 | 3.43E-01 | 5.37E-01 | 7.06E-01 | 3.16E-01 | 9.87E-01 | 8.38E-01 | 2.80E-01 | 6.86E-01 |
| sex | 3.39E-01 | 2.36E-01 | 2.21E-01 | 2.39E-01 | 1.53E-01 | 1.71E-01 | 3.98E-01 | 2.08E-01 | 2.62E-01 | 7.45E-01 | 3.57E-01 | 6.61E-01 | **3.56E-02** | **3.68E-02** | **2.93E-02** | **4.57E-02** |
| apoe4 | **1.65E-03** | **1.10E-04** | 5.80E-01 | **8.30E-03** | **2.81E-02** | **1.22E-03** | 7.22E-01 | **2.64E-02** | 9.06E-02 | **1.58E-04** | 5.11E-01 | **8.67E-03** | 1.01E-01 | **1.23E-03** | 4.32E-01 | 1.71E-01 |
| raceB | 8.50E-01 | 5.86E-01 | 9.08E-01 | 8.29E-01 | 9.62E-01 | 4.48E-01 | 5.57E-01 | 5.89E-01 | 9.25E-01 | 9.02E-01 | 7.13E-01 | 9.20E-01 | - | - | - | - |
| raceH | 4.24E-01 | 6.92E-01 | 7.01E-01 | 5.13E-01 | 5.67E-01 | 9.55E-01 | 9.49E-01 | 7.32E-01 | 6.24E-01 | 6.20E-01 | 8.10E-01 | 5.73E-01 | 8.43E-02 | **2.50E-02** | 3.05E-01 | **4.90E-02** |
| raceW | 8.37E-01 | 4.79E-01 | 6.83E-01 | 7.79E-01 | 9.02E-01 | 4.33E-01 | 4.98E-01 | 5.88E-01 | 9.02E-01 | 7.56E-01 | 6.86E-01 | 8.38E-01 | 6.73E-01 | 3.55E-01 | 9.60E-02 | 5.96E-01 |
| RIN | 7.37E-01 | 3.80E-01 | 3.34E-01 | 7.02E-01 | 4.75E-01 | 4.88E-01 | 3.04E-01 | 8.85E-01 | 6.69E-01 | 5.61E-01 | 1.77E-01 | 7.72E-01 | 7.59E-02 | 2.12E-01 | **3.88E-02** | 3.41E-01 |
| PMI | 3.34E-01 | **9.29E-03** | **3.25E-03** | **1.61E-02** | 6.18E-01 | 5.88E-02 | **1.93E-03** | 1.14E-01 | 8.18E-01 | **4.71E-02** | **4.13E-02** | 1.02E-01 | 5.70E-01 | 6.06E-02 | **8.55E-03** | 6.04E-02 |
| Multiple R^2 | 0.31 | 0.25 | 0.30 | 0.35 | 0.26 | 0.23 | 0.26 | 0.30 | 0.27 | 0.23 | 0.34 | 0.39 | 0.31 | 0.35 | 0.27 | 0.35 |
| Adjusted R^2 | 0.26 | 0.20 | 0.24 | 0.30 | 0.20 | 0.16 | 0.19 | 0.24 | 0.19 | 0.15 | 0.27 | 0.32 | 0.26 | 0.30 | 0.21 | 0.30 |

**Table S7 Index genes and their weights contributing to the deep learning model.**

| ensembl ID | symbol | weight | module |
|---|---|---|---|
| ENSG00000088836 | SLC4A11 | 1 | green |
| ENSG00000131095 | GFAP | 0.8831 | blue |
| ENSG00000274276 | CBSL | 0.76773 | grey |
| ENSG00000196517 | SLC6A9 | 0.75606 | green |
| ENSG00000158169 | FANCC | 0.7318 | green |
| ENSG00000168743 | NPNT | 0.66154 | turquoise |
| ENSG00000125337 | KIF25 | 0.63187 | grey |
| ENSG00000262877 | AC110285.2 | 0.6311 | grey |
| ENSG00000134533 | RERG | 0.62733 | blue |
| ENSG00000153822 | KCNJ16 | 0.57374 | blue |
| ENSG00000183090 | FREM3 | 0.55655 | turquoise |
| ENSG00000140368 | PSTPIP1 | 0.52183 | blue |
| ENSG00000182648 | LINC01006 | 0.50898 | turquoise |
| ENSG00000196415 | PRTN3 | 0.50515 | turquoise |
| ENSG00000115602 | IL1RL1 | 0.43866 | brown |
| ENSG00000205611 | LINC01597 | 0.43353 | turquoise |
| ENSG00000140678 | ITGAX | 0.40071 | brown |
| ENSG00000189343 | RPS2P46 | 0.39502 | turquoise |
| ENSG00000225302 |  | 0.38756 | grey |
| ENSG00000275830 | AL355974.2 | 0.38418 | brown |
| ENSG00000169031 | COL4A3 | 0.36828 | brown |
| ENSG00000166012 | TAF1D | 0.36241 | turquoise |
| ENSG00000020577 | SAMD4A | 0.36187 | brown |
| ENSG00000154864 | PIEZO2 | 0.35731 | green |
| ENSG00000105643 | ARRDC2 | 0.34802 | brown |
| ENSG00000206195 | DUXAP8 | 0.33919 | grey |
| ENSG00000111788 | AC009533.1 | 0.33254 | grey |
| ENSG00000136297 | MMD2 | 0.30094 | blue |
| ENSG00000253159 | PCDHGA12 | 0.29011 | blue |
| ENSG00000163319 | MRPS18C | 0.28927 | turquoise |
| ENSG00000138778 | CENPE | 0.2886 | yellow |
| ENSG00000139410 | SDSL | 0.28818 | turquoise |
| ENSG00000148655 | LRMDA | 0.28607 | grey |
| ENSG00000228314 | CYP4F29P | 0.28601 | grey |
| ENSG00000136449 | MYCBPAP | 0.28394 | turquoise |
| ENSG00000198830 | HMGN2 | 0.28372 | turquoise |
| ENSG00000142910 | TINAGL1 | 0.27878 | brown |
| ENSG00000146250 | PRSS35 | 0.27569 | blue |
| ENSG00000157005 | SST | 0.26298 | turquoise |
| ENSG00000151233 | GXYLT1 | 0.25542 | blue |
| ENSG00000125534 | PPDPF | 0.24935 | turquoise |
| ENSG00000136732 | GYPC | 0.24319 | brown |
| ENSG00000176387 | HSD11B2 | 0.24242 | grey |
| ENSG00000143546 | S100A8 | 0.24016 | brown |
| ENSG00000246379 | AC007495.1 | 0.23992 | yellow |
| ENSG00000279267 | AL078621.3 | 0.23269 | turquoise |
| ENSG00000184730 | APOBR | 0.23222 | brown |
| ENSG00000172572 | PDE3A | 0.23215 | blue |
| ENSG00000105419 | MEIS3 | 0.22922 | turquoise |
| ENSG00000265688 | MAFG-AS1 | 0.22683 | yellow |
| ENSG00000144837 | PLA1A | 0.22175 | brown |
| ENSG00000229807 | XIST | 0.21942 | grey |
| ENSG00000236963 | LINC01141 | 0.21719 | turquoise |
| ENSG00000177464 | GPR4 | 0.21254 | brown |
| ENSG00000139971 | C14orf37 | 0.21127 | turquoise |
| ENSG00000196436 | NPIPB15 | 0.20216 | grey |
| ENSG00000237438 | CECR7 | 0.19829 | turquoise |
| ENSG00000232528 | AL109809.1 | 0.1953 | yellow |
| ENSG00000081052 | COL4A4 | 0.19407 | brown |
| ENSG00000236819 | LINC01563 | 0.19138 | turquoise |
| ENSG00000246982 | Z84485.1 | 0.18968 | grey |
| ENSG00000162992 | NEUROD1 | 0.18426 | turquoise |
| ENSG00000166573 | GALR1 | 0.18271 | turquoise |
| ENSG00000240583 | AQP1 | 0.17767 | blue |
| ENSG00000269707 | AC018730.2 | 0.17621 | turquoise |
| ENSG00000140961 | OSGIN1 | 0.17176 | grey |
| ENSG00000196812 | ZSCAN16 | 0.17081 | turquoise |
| ENSG00000204128 | C2orf72 | 0.17066 | turquoise |
| ENSG00000244734 | HBB | 0.1691 | grey |
| ENSG00000141665 | FBXO15 | 0.16457 | turquoise |
| ENSG00000070731 | ST6GALNAC2 | 0.16349 | brown |
| ENSG00000118515 | SGK1 | 0.16158 | green |
| ENSG00000111181 | SLC6A12 | 0.16093 | green |
| ENSG00000013573 | DDX11 | 0.16032 | grey |
| ENSG00000163220 | S100A9 | 0.15697 | brown |
| ENSG00000183876 | ARSI | 0.15489 | blue |
| ENSG00000102287 | GABRE | 0.15444 | brown |
| ENSG00000099260 | PALMD | 0.15344 | brown |
| ENSG00000233695 | GAS6-AS1 | 0.14852 | turquoise |
| ENSG00000143429 | AC116050.1 | 0.14798 | yellow |
| ENSG00000162551 | ALPL | 0.147 | brown |
| ENSG00000125810 | CD93 | 0.1433 | brown |
| ENSG00000100376 | FAM118A | 0.1433 | grey |
| ENSG00000240342 | RPS2P5 | 0.14158 | grey |
| ENSG00000226067 | LINC00623 | 0.14089 | turquoise |
| ENSG00000136982 | DSCC1 | 0.14021 | turquoise |
| ENSG00000155980 | KIF5A | 0.14008 | blue |
| ENSG00000198624 | CCDC69 | 0.13866 | green |
| ENSG00000143318 | CASQ1 | 0.13852 | turquoise |
| ENSG00000176809 | LRRC37A3 | 0.13851 | turquoise |
| ENSG00000006611 | USH1C | 0.13791 | blue |
| ENSG00000138336 | TET1 | 0.13733 | blue |
| ENSG00000110492 | MDK | 0.13713 | turquoise |
| ENSG00000149418 | ST14 | 0.13659 | turquoise |
| ENSG00000056998 | GYG2 | 0.13583 | blue |
| ENSG00000198554 | WDHD1 | 0.13521 | turquoise |
| ENSG00000261609 | GAN | 0.13335 | turquoise |
| ENSG00000224383 | PRR29 | 0.13308 | brown |
| ENSG00000126709 | IFI6 | 0.13304 | turquoise |
| ENSG00000154734 | ADAMTS1 | 0.13281 | brown |
| ENSG00000133466 | C1QTNF6 | 0.13189 | grey |
| ENSG00000272602 | ZNF595 | 0.13104 | turquoise |
| ENSG00000135269 | TES | 0.13042 | brown |
| ENSG00000147509 | RGS20 | 0.12943 | blue |
| ENSG00000223865 | HLA-DPB1 | 0.12734 | brown |
| ENSG00000157379 | DHRS1 | 0.12687 | turquoise |
| ENSG00000260942 | CAPN10-AS1 | 0.12653 | grey |
| ENSG00000205517 | RGL3 | 0.12633 | brown |
| ENSG00000127325 | BEST3 | 0.12603 | blue |
| ENSG00000272690 | LINC02018 | 0.12532 | grey |
| ENSG00000160200 | CBS | 0.12513 | grey |
| ENSG00000181019 | NQO1 | 0.12456 | blue |
| ENSG00000231768 | LINC01354 | 0.12396 | blue |
| ENSG00000188848 | BEND4 | 0.12376 | turquoise |
| ENSG00000146215 | CRIP3 | 0.12269 | turquoise |
| ENSG00000112964 | GHR | 0.12121 | turquoise |
| ENSG00000162493 | PDPN | 0.11911 | blue |
| ENSG00000140522 | RLBP1 | 0.11496 | blue |
| ENSG00000271254 | AC240274.1 | 0.11495 | turquoise |
| ENSG00000215559 | ANKRD20A11P | 0.11436 | yellow |
| ENSG00000132832 | AL139352.1 | 0.11323 | turquoise |
| ENSG00000170379 | TCAF2 | 0.11205 | grey |

| ensembl ID | symbol | weight | module |
|---|---|---|---|
| ENSG00000267568 | AC016168.2 | 0.11183 | grey |
| ENSG00000242686 | AC107464.1 | 0.11139 | turquoise |
| ENSG00000114656 | KIAA1257 | 0.11108 | yellow |
| ENSG00000234327 | AC012146.1 | 0.11097 | turquoise |
| ENSG00000136859 | ANGPTL2 | 0.11076 | green |
| ENSG00000118523 | CTGF | 0.10811 | brown |
| ENSG00000255769 | GOLGA2P10 | 0.1075 | turquoise |
| ENSG00000130287 | NCAN | 0.10637 | blue |
| ENSG00000185864 | NPIPB4 | 0.10473 | turquoise |
| ENSG00000140807 | NKD1 | 0.10394 | green |
| ENSG00000188916 | FAM196A | 0.10389 | turquoise |
| ENSG00000179477 | ALOX12B | 0.10367 | turquoise |
| ENSG00000118777 | ABCG2 | 0.10156 | brown |
| ENSG00000205336 | ADGRG1 | 0.10133 | blue |
| ENSG00000087116 | ADAMTS2 | 0.10058 | turquoise |
| ENSG00000136826 | KLF4 | 0.09994 | brown |
| ENSG00000123454 | DBH | 0.099888 | turquoise |
| ENSG00000137404 | NRM | 0.098639 | brown |
| ENSG00000182109 | | 0.098368 | turquoise |
| ENSG00000121281 | ADCY7 | 0.097449 | turquoise |
| ENSG00000251442 | LINC01094 | 0.096568 | brown |
| ENSG00000273702 | AC091271.1 | 0.096279 | blue |
| ENSG00000107821 | KAZALD1 | 0.095312 | turquoise |
| ENSG00000129204 | USP6 | 0.093962 | turquoise |
| ENSG00000243244 | STON1 | 0.093914 | brown |
| ENSG00000233251 | AC007743.1 | 0.093808 | turquoise |
| ENSG00000270231 | NBPF8 | 0.093277 | turquoise |
| ENSG00000126249 | PDCD2L | 0.092782 | turquoise |
| ENSG00000188039 | NWD1 | 0.092777 | green |
| ENSG00000119509 | INVS | 0.092774 | turquoise |
| ENSG00000121858 | TNFSF10 | 0.092424 | grey |
| ENSG00000006534 | ALDH3B1 | 0.090819 | blue |
| ENSG00000099958 | DERL3 | 0.090165 | grey |
| ENSG00000104899 | AMH | 0.089358 | grey |
| ENSG00000112210 | RAB23 | 0.088158 | turquoise |
| ENSG00000244026 | FAM86DP | 0.087718 | turquoise |
| ENSG00000181631 | P2RY13 | 0.087613 | grey |
| ENSG00000112149 | CD83 | 0.087075 | turquoise |
| ENSG00000281501 | SEPSECS-AS1 | 0.086068 | yellow |
| ENSG00000254726 | MEX3A | 0.085666 | grey |
| ENSG00000158406 | HIST1H4H | 0.08508 | grey |
| ENSG00000227953 | LINC01341 | 0.083835 | turquoise |
| ENSG00000103742 | IGDCC4 | 0.08377 | blue |
| ENSG00000105707 | HPN | 0.083057 | green |
| ENSG00000133069 | TMCC2 | 0.082881 | turquoise |
| ENSG00000156076 | WIF1 | 0.0828 | blue |
| ENSG00000198502 | HLA-DRB5 | 0.082379 | brown |
| ENSG00000232931 | LINC00342 | 0.081125 | turquoise |
| ENSG00000178796 | RIIAD1 | 0.080697 | turquoise |
| ENSG00000180481 | GLIPR1L2 | 0.080023 | grey |
| ENSG00000233058 | LINC00884 | 0.079546 | turquoise |
| ENSG00000171243 | SOSTDC1 | 0.079334 | turquoise |
| ENSG00000141179 | PCTP | 0.079217 | turquoise |
| ENSG00000211448 | DIO2 | 0.078864 | grey |
| ENSG00000142621 | FHAD1 | 0.078678 | turquoise |
| ENSG00000163520 | FBLN2 | 0.078269 | turquoise |
| ENSG00000116830 | TTF2 | 0.077922 | turquoise |
| ENSG00000173918 | C1QTNF1 | 0.077794 | brown |
| ENSG00000182511 | FES | 0.076993 | brown |
| ENSG00000175164 | ABO | 0.076358 | brown |
| ENSG00000227827 | AC138969.2 | 0.076133 | turquoise |
| ENSG00000169246 | NPIPB3 | 0.075304 | turquoise |
| ENSG00000235072 | AC012074.1 | 0.074836 | turquoise |

| ensembl ID | symbol | weight | module |
|---|---|---|---|
| ENSG00000273142 | AC073335.2 | 0.073886 | grey |
| ENSG00000149150 | SLC43A1 | 0.073885 | turquoise |
| ENSG00000054392 | HHAT | 0.073682 | turquoise |
| ENSG00000277494 | GPIHBP1 | 0.072773 | green |
| ENSG00000224914 | LINC00863 | 0.072222 | turquoise |
| ENSG00000260948 | AL390195.2 | 0.07217 | turquoise |
| ENSG00000186918 | ZNF395 | 0.072135 | blue |
| ENSG00000144057 | ST6GAL2 | 0.071791 | turquoise |
| ENSG00000162383 | SLC1A7 | 0.071448 | brown |
| ENSG00000162836 | ACP6 | 0.070934 | blue |
| ENSG00000188681 | TEKT4P2 | 0.068776 | grey |
| ENSG00000214425 | LRRC37A4P | 0.06871 | yellow |
| ENSG00000173947 | PIFO | 0.068495 | blue |
| ENSG00000072736 | NFATC3 | 0.068112 | turquoise |
| ENSG00000111962 | UST | 0.067421 | turquoise |
| ENSG00000244480 | AC005154.3 | 0.067129 | turquoise |
| ENSG00000196154 | S100A4 | 0.066933 | green |
| ENSG00000148357 | HMCN2 | 0.066764 | turquoise |
| ENSG00000134323 | MYCN | 0.066344 | turquoise |
| ENSG00000169418 | NPR1 | 0.065934 | brown |
| ENSG00000277758 | FO681492.1 | 0.065003 | turquoise |
| ENSG00000105675 | ATP4A | 0.064717 | turquoise |
| ENSG00000267838 | AC245884.8 | 0.06406 | turquoise |
| ENSG00000141469 | SLC14A1 | 0.061732 | blue |
| ENSG00000255545 | AP004608.1 | 0.061171 | turquoise |
| ENSG00000006047 | YBX2 | 0.060636 | turquoise |
| ENSG00000185519 | FAM131C | 0.059725 | turquoise |
| ENSG00000260426 | AC008060.4 | 0.059616 | turquoise |
| ENSG00000139675 | HNRNPA1L2 | 0.059582 | turquoise |
| ENSG00000196196 | HRCT1 | 0.058657 | grey |
| ENSG00000177551 | NHLH2 | 0.058474 | turquoise |
| ENSG00000077585 | GPR137B | 0.057915 | blue |
| ENSG00000228716 | DHFR | 0.05731 | grey |
| ENSG00000157833 | GAREM2 | 0.056805 | green |
| ENSG00000176641 | RNF152 | 0.056721 | brown |
| ENSG00000177425 | PAWR | 0.056696 | brown |
| ENSG00000011426 | ANLN | 0.055581 | green |
| ENSG00000016602 | CLCA4 | 0.05436 | green |
| ENSG00000173801 | JUP | 0.05383 | blue |
| ENSG00000250903 | GMDS-AS1 | 0.053526 | turquoise |
| ENSG00000149809 | TM7SF2 | 0.051891 | turquoise |
| ENSG00000188536 | HBA2 | 0.050951 | yellow |
| ENSG00000144401 | METTL21A | 0.050115 | turquoise |
| ENSG00000280670 | CCDC163 | 0.049977 | blue |
| ENSG00000196369 | SRGAP2B | 0.049672 | turquoise |
| ENSG00000175643 | RMI2 | 0.048871 | turquoise |
| ENSG00000166473 | PKD1L2 | 0.048246 | green |
| ENSG00000081853 | PCDHGA2 | 0.048241 | blue |
| ENSG00000152527 | PLEKHH2 | 0.047905 | green |
| ENSG00000174807 | CD248 | 0.047572 | brown |
| ENSG00000171365 | CLCN5 | 0.047279 | turquoise |
| ENSG00000203883 | SOX18 | 0.047181 | yellow |
| ENSG00000242299 | AC073861.1 | 0.046946 | turquoise |
| ENSG00000253710 | ALG11 | 0.046563 | grey |
| ENSG00000117115 | PADI2 | 0.046323 | green |
| ENSG00000087245 | MMP2 | 0.046031 | brown |
| ENSG00000177076 | ACER2 | 0.045974 | turquoise |
| ENSG00000154764 | WNT7A | 0.045971 | turquoise |
| ENSG00000179954 | SSC5D | 0.045913 | turquoise |
| ENSG00000003400 | CASP10 | 0.045252 | brown |
| ENSG00000140545 | MFGE8 | 0.044562 | turquoise |
| ENSG00000167840 | ZNF232 | 0.0444 | turquoise |
| ENSG00000145014 | TMEM44 | 0.04418 | turquoise |

| ensembl ID | symbol | weight | module |
| --- | --- | --- | --- |
| ENSG00000120903 | CHRNA2 | 0.04401 | turquoise |
| ENSG00000122085 | MTERF4 | 0.043004 | turquoise |
| ENSG00000183379 | SYNDIG1L | 0.042898 | turquoise |
| ENSG00000064666 | CNN2 | 0.042205 | brown |
| ENSG00000116711 | PLA2G4A | 0.04198 | turquoise |
| ENSG00000241015 | TPM3P9 | 0.041974 | turquoise |
| ENSG00000225968 | ELFN1 | 0.041724 | turquoise |
| ENSG00000186960 | LINC01551 | 0.041551 | turquoise |
| ENSG00000225313 | AL513327.1 | 0.041533 | turquoise |
| ENSG00000270015 | AC087481.3 | 0.041075 | turquoise |
| ENSG00000165084 | C8orf34 | 0.040337 | turquoise |
| ENSG00000122557 | HERPUD2 | 0.040103 | turquoise |
| ENSG00000215440 | NPEPL1 | 0.039073 | grey |
| ENSG00000183506 | PI4KAP2 | 0.039042 | turquoise |
| ENSG00000201136 | RNU6-353P | 0.038874 | yellow |
| ENSG00000151240 | DIP2C | 0.038573 | turquoise |
| ENSG00000155254 | MARVELD1 | 0.038509 | turquoise |
| ENSG00000198821 | CD247 | 0.038257 | turquoise |
| ENSG00000103226 | NOMO3 | 0.037798 | turquoise |
| ENSG00000186231 | KLHL32 | 0.037312 | turquoise |
| ENSG00000122375 | OPN4 | 0.037269 | turquoise |
| ENSG00000196132 | MYT1 | 0.037256 | green |
| ENSG00000214756 | CSKMT | 0.036588 | turquoise |
| ENSG00000166483 | WEE1 | 0.036321 | blue |
| ENSG00000104332 | SFRP1 | 0.036097 | turquoise |
| ENSG00000205918 | PDPK2P | 0.035848 | turquoise |
| ENSG00000130055 | GDPD2 | 0.035549 | blue |
| ENSG00000271743 | AF287957.1 | 0.035312 | turquoise |
| ENSG00000125378 | BMP4 | 0.03498 | turquoise |
| ENSG00000115339 | GALNT3 | 0.034953 | turquoise |
| ENSG00000135097 | MSI1 | 0.034616 | blue |
| ENSG00000107562 | CXCL12 | 0.034598 | grey |
| ENSG00000261819 | AC138932.3 | 0.034368 | yellow |
| ENSG00000152208 | GRID2 | 0.034281 | grey |
| ENSG00000183496 | MEX3B | 0.034195 | turquoise |
| ENSG00000110628 | SLC22A18 | 0.033648 | turquoise |
| ENSG00000172548 | NIPAL4 | 0.033169 | green |
| ENSG00000154721 | JAM2 | 0.032771 | blue |
| ENSG00000277702 | AC239859.6 | 0.032637 | turquoise |
| ENSG00000116874 | WARS2 | 0.03253 | turquoise |
| ENSG00000213096 | ZNF254 | 0.032354 | turquoise |
| ENSG00000134253 | TRIM45 | 0.031874 | turquoise |
| ENSG00000188234 | AGAP4 | 0.031704 | turquoise |
| ENSG00000225178 | RPSAP58 | 0.031111 | grey |
| ENSG00000250802 | ZBED3-AS1 | 0.031101 | blue |
| ENSG00000256269 | HMBS | 0.030967 | turquoise |
| ENSG00000133519 | ZDHHC8P1 | 0.030901 | turquoise |
| ENSG00000230177 | AL080317.1 | 0.030673 | turquoise |
| ENSG00000079462 | PAFAH1B3 | 0.030456 | turquoise |
| ENSG00000227191 | TRGC2 | 0.030352 | grey |
| ENSG00000101883 | RHOXF1 | 0.030191 | grey |
| ENSG00000245910 | SNHG6 | 0.029881 | turquoise |
| ENSG00000259429 | UBE2Q2P2 | 0.029878 | yellow |
| ENSG00000179029 | TMEM107 | 0.029857 | turquoise |
| ENSG00000064300 | NGFR | 0.029709 | brown |
| ENSG00000255198 | SNHG9 | 0.029379 | yellow |
| ENSG00000015133 | CCDC88C | 0.029173 | turquoise |
| ENSG00000230202 | AL450405.1 | 0.029077 | yellow |
| ENSG00000224195 | AC022400.1 | 0.028376 | turquoise |
| ENSG00000215374 | FAM66B | 0.028089 | turquoise |
| ENSG00000188290 | HES4 | 0.02764 | yellow |
| ENSG00000186891 | TNFRSF18 | 0.027351 | turquoise |
| ENSG00000282936 | AC004706.4 | 0.027141 | turquoise |

| ensembl ID | symbol | weight | module |
| --- | --- | --- | --- |
| ENSG00000102882 | MAPK3 | 0.026837 | turquoise |
| ENSG00000169908 | TM4SF1 | 0.026577 | brown |
| ENSG00000214189 | ZNF788 | 0.026498 | blue |
| ENSG00000167123 | CERCAM | 0.026382 | green |
| ENSG00000171806 | METTL18 | 0.02636 | turquoise |
| ENSG00000272010 | AC100814.1 | 0.026355 | turquoise |
| ENSG00000180834 | MAP6D1 | 0.026039 | turquoise |
| ENSG00000229036 | VDAC1P8 | 0.025499 | turquoise |
| ENSG00000198826 | ARHGAP11A | 0.0252 | turquoise |
| ENSG00000140750 | ARHGAP17 | 0.025171 | green |
| ENSG00000101224 | CDC25B | 0.024869 | turquoise |
| ENSG00000149531 | FRG1BP | 0.024799 | turquoise |
| ENSG00000157613 | CREB3L1 | 0.024696 | turquoise |
| ENSG00000095383 | TBC1D2 | 0.024379 | green |
| ENSG00000049283 | EPN3 | 0.023883 | turquoise |
| ENSG00000165092 | ALDH1A1 | 0.02377 | turquoise |
| ENSG00000113209 | PCDHB5 | 0.023537 | blue |
| ENSG00000225630 | MTND2P28 | 0.023532 | turquoise |
| ENSG00000255020 | AF131216.3 | 0.023446 | turquoise |
| ENSG00000131477 | RAMP2 | 0.023316 | grey |
| ENSG00000134899 | ERCC5 | 0.023211 | grey |
| ENSG00000127528 | KLF2 | 0.023196 | yellow |
| ENSG00000129993 | CBFA2T3 | 0.022885 | turquoise |
| ENSG00000071894 | CPSF1 | 0.022546 | turquoise |
| ENSG00000132275 | RRP8 | 0.022522 | turquoise |
| ENSG00000204172 | AGAP9 | 0.02244 | turquoise |
| ENSG00000162976 | PQLC3 | 0.022311 | turquoise |
| ENSG00000183091 | NEB | 0.021864 | turquoise |
| ENSG00000167720 | SRR | 0.021825 | turquoise |
| ENSG00000233901 | LINC01503 | 0.021384 | turquoise |
| ENSG00000272971 | AL365181.4 | 0.021305 | turquoise |
| ENSG00000105750 | ZNF85 | 0.021183 | turquoise |
| ENSG00000198547 | C20orf203 | 0.020848 | turquoise |
| ENSG00000105287 | PRKD2 | 0.02078 | grey |
| ENSG00000273748 | AL592183.1 | 0.020534 | grey |
| ENSG00000108375 | RNF43 | 0.02045 | blue |
| ENSG00000154262 | ABCA6 | 0.020446 | yellow |
| ENSG00000187678 | SPRY4 | 0.020403 | turquoise |
| ENSG00000277053 | GTF2IP1 | 0.020253 | turquoise |
| ENSG00000258702 | AL137786.1 | 0.020096 | turquoise |
| ENSG00000215769 | ARHGAP27P1-BPTFP1-KPNA2P3 | 0.019563 | turquoise |
| ENSG00000280187 | AC022107.1 | 0.019505 | turquoise |
| ENSG00000134207 | SYT6 | 0.019458 | turquoise |
| ENSG00000197993 | KEL | 0.019368 | green |
| ENSG00000141314 | RHBDL3 | 0.019215 | turquoise |
| ENSG00000172301 | COPRS | 0.019208 | turquoise |
| ENSG00000188242 | AC010442.1 | 0.019108 | turquoise |
| ENSG00000258010 | AC016705.1 | 0.018708 | turquoise |
| ENSG00000120071 | KANSL1 | 0.01824 | turquoise |
| ENSG00000261126 | RBFADN | 0.017896 | turquoise |
| ENSG00000162745 | OLFML2B | 0.01788 | turquoise |
| ENSG00000115318 | LOXL3 | 0.017866 | blue |
| ENSG00000280255 | AC004947.2 | 0.01785 | turquoise |
| ENSG00000177335 | C8orf31 | 0.0178 | turquoise |
| ENSG00000134020 | PEBP4 | 0.017625 | turquoise |
| ENSG00000261377 | PDCD6IPP2 | 0.017538 | turquoise |
| ENSG00000157978 | LDLRAP1 | 0.017277 | green |
| ENSG00000272419 | AC241585.2 | 0.017195 | turquoise |
| ENSG00000155792 | DEPTOR | 0.017184 | turquoise |
| ENSG00000153495 | TEX29 | 0.017141 | turquoise |

| ensembl ID | symbol | weight | module |
|---|---|---|---|
| ENSG00000129534 | MIS18BP1 | 0.016773 | turquoise |
| ENSG00000066294 | CD84 | 0.016549 | brown |
| ENSG00000261386 | AC027682.4 | 0.016451 | turquoise |
| ENSG00000183921 | SDR42E2 | 0.016247 | turquoise |
| ENSG00000259495 | AC016705.2 | 0.016217 | turquoise |
| ENSG00000169857 | AVEN | 0.016122 | turquoise |
| ENSG00000135046 | ANXA1 | 0.016084 | brown |
| ENSG00000139352 | ASCL1 | 0.015893 | green |
| ENSG00000103995 | CEP152 | 0.015656 | yellow |
| ENSG00000204611 | ZNF616 | 0.015592 | turquoise |
| ENSG00000154263 | ABCA10 | 0.015493 | turquoise |
| ENSG00000242732 | RTL5 | 0.01514 | turquoise |
| ENSG00000088053 | GP6 | 0.014677 | turquoise |
| ENSG00000136319 | TTC5 | 0.014085 | turquoise |
| ENSG00000215915 | ATAD3C | 0.013768 | turquoise |
| ENSG00000161940 | BCL6B | 0.013534 | brown |
| ENSG00000173163 | COMMD1 | 0.013398 | turquoise |
| ENSG00000222044 | AL031587.1 | 0.013157 | yellow |
| ENSG00000261353 | | 0.012984 | grey |
| ENSG00000280351 | AC127496.7 | 0.012956 | yellow |
| ENSG00000204323 | SMIM5 | 0.012679 | green |
| ENSG00000078295 | ADCY2 | 0.012572 | blue |
| ENSG00000125850 | OVOL2 | 0.012543 | turquoise |
| ENSG00000272668 | AL590560.2 | 0.012513 | grey |
| ENSG00000147571 | CRH | 0.012283 | turquoise |
| ENSG00000206344 | HCG27 | 0.012264 | yellow |
| ENSG00000185269 | NOTUM | 0.012222 | turquoise |
| ENSG00000112773 | FAM46A | 0.012208 | brown |
| ENSG00000138942 | RNF185 | 0.011899 | turquoise |
| ENSG00000268751 | SCGB1B2P | 0.011876 | yellow |
| ENSG00000242220 | TCP10L | 0.011647 | turquoise |
| ENSG00000173653 | RCE1 | 0.011606 | turquoise |
| ENSG00000279656 | AL132780.4 | 0.011584 | grey |
| ENSG00000164904 | ALDH7A1 | 0.011286 | blue |
| ENSG00000064787 | BCAS1 | 0.011277 | turquoise |
| ENSG00000261373 | VPS9D1-AS1 | 0.011161 | yellow |
| ENSG00000101871 | MID1 | 0.011119 | blue |
| ENSG00000248049 | UBA6-AS1 | 0.011014 | turquoise |
| ENSG00000269935 | AC092720.2 | 0.010962 | turquoise |
| ENSG00000123689 | G0S2 | 0.010793 | grey |
| ENSG00000273308 | AC024560.3 | 0.010598 | yellow |
| ENSG00000138741 | TRPC3 | 0.010503 | turquoise |
| ENSG00000165553 | NGB | 0.010303 | turquoise |
| ENSG00000266338 | NBPF15 | 0.010285 | turquoise |
| ENSG00000132837 | DMGDH | 0.010225 | turquoise |
| ENSG00000254081 | LINC01299 | 0.010091 | grey |
| ENSG00000065268 | WDR18 | 0.0099383 | turquoise |
| ENSG00000133433 | GSTT2B | 0.0097166 | turquoise |
| ENSG00000114779 | ABHD14B | 0.0094285 | turquoise |
| ENSG00000220804 | LINC01881 | 0.0093006 | yellow |
| ENSG00000101282 | RSPO4 | 0.0089581 | turquoise |
| ENSG00000243742 | RPLP0P2 | 0.0086551 | grey |
| ENSG00000144283 | PKP4 | 0.0081785 | turquoise |
| ENSG00000111364 | DDX55 | 0.0078924 | yellow |
| ENSG00000185904 | LINC00839 | 0.0078883 | turquoise |
| ENSG00000136114 | THSD1 | 0.0078444 | blue |
| ENSG00000220008 | LINGO3 | 0.0077896 | yellow |
| ENSG00000227782 | AC002553.1 | 0.0075723 | yellow |
| ENSG00000118514 | ALDH8A1 | 0.0074187 | turquoise |
| ENSG00000203867 | RBM20 | 0.007079 | turquoise |
| ENSG00000108370 | RGS9 | 0.006996 | blue |
| ENSG00000143994 | ABHD1 | 0.0069629 | blue |
| ENSG00000251429 | AC098679.2 | 0.0069347 | green |
| ENSG00000101311 | FERMT1 | 0.0069089 | green |
| ENSG00000167874 | TMEM88 | 0.0068412 | grey |
| ENSG00000075461 | CACNG4 | 0.0067956 | green |
| ENSG00000164818 | DNAAF5 | 0.0067319 | turquoise |
| ENSG00000181773 | GPR3 | 0.0066865 | turquoise |
| ENSG00000136147 | PHF11 | 0.0066154 | turquoise |
| ENSG00000267280 | TBX2-AS1 | 0.006453 | brown |
| ENSG00000198938 | MT-CO3 | 0.0064236 | turquoise |
| ENSG00000120279 | MYCT1 | 0.0063932 | brown |
| ENSG00000160999 | SH2B2 | 0.0063499 | yellow |
| ENSG00000260025 | AC009414.2 | 0.006246 | turquoise |
| ENSG00000280485 | AL358154.1 | 0.0061272 | yellow |
| ENSG00000182853 | VMO1 | 0.0060622 | turquoise |
| ENSG00000140795 | MYLK3 | 0.006018 | turquoise |
| ENSG00000173457 | PPP1R14B | 0.0058658 | turquoise |
| ENSG00000172985 | SH3RF3 | 0.005857 | turquoise |
| ENSG00000214491 | SEC14L6 | 0.0058494 | turquoise |
| ENSG00000103485 | QPRT | 0.0058033 | turquoise |
| ENSG00000028116 | VRK2 | 0.0055534 | green |
| ENSG00000171094 | ALK | 0.0052428 | turquoise |
| ENSG00000197536 | C5orf56 | 0.0052398 | turquoise |
| ENSG00000272674 | PCDHB16 | 0.0051392 | turquoise |
| ENSG00000270728 | AL035413.2 | 0.0051353 | yellow |
| ENSG00000144061 | NPHP1 | 0.004951 | turquoise |
| ENSG00000177984 | LCN15 | 0.004577 | turquoise |
| ENSG00000102390 | PBDC1 | 0.0045051 | turquoise |
| ENSG00000132793 | LPIN3 | 0.0043651 | blue |
| ENSG00000255836 | AC131206.1 | 0.0042159 | yellow |
| ENSG00000166086 | JAM3 | 0.0041743 | turquoise |
| ENSG00000272512 | AL645608.8 | 0.0041256 | grey |
| ENSG00000238197 | PAXBP1-AS1 | 0.0039422 | turquoise |
| ENSG00000135824 | RGS8 | 0.0037904 | turquoise |
| ENSG00000163485 | ADORA1 | 0.0037369 | turquoise |
| ENSG00000175768 | TOMM5 | 0.0035518 | turquoise |
| ENSG00000280734 | LINC01232 | 0.0034469 | turquoise |
| ENSG00000170684 | ZNF296 | 0.0034448 | yellow |
| ENSG00000132846 | ZBED3 | 0.0034333 | green |
| ENSG00000198468 | FLVCR1-AS1 | 0.0032206 | grey |
| ENSG00000183066 | WBP2NL | 0.0030944 | yellow |
| ENSG00000137269 | LRRC1 | 0.0030549 | green |
| ENSG00000169064 | ZBBX | 0.0029782 | turquoise |
| ENSG00000198727 | MT-CYB | 0.002951 | turquoise |
| ENSG00000140022 | STON2 | 0.0028454 | blue |
| ENSG00000196502 | SULT1A1 | 0.0028402 | turquoise |
| ENSG00000260912 | AL158206.1 | 0.0028194 | yellow |
| ENSG00000267107 | PCAT19 | 0.0028068 | brown |
| ENSG00000170011 | MYRIP | 0.0027228 | turquoise |
| ENSG00000196735 | HLA-DQA1 | 0.0026605 | brown |
| ENSG00000198886 | MT-ND4 | 0.0026325 | turquoise |
| ENSG00000196418 | ZNF124 | 0.0026246 | yellow |
| ENSG00000205744 | DENND1C | 0.0026099 | brown |
| ENSG00000137766 | UNC13C | 0.0026098 | turquoise |
| ENSG00000125804 | FAM182A | 0.0025733 | grey |
| ENSG00000146834 | MEPCE | 0.0025464 | turquoise |
| ENSG00000180354 | MTURN | 0.002539 | turquoise |
| ENSG00000171388 | APLN | 0.0024605 | green |
| ENSG00000228543 | AC003684.1 | 0.0024508 | grey |
| ENSG00000187775 | DNAH17 | 0.0024143 | green |
| ENSG00000152133 | GPATCH11 | 0.0023659 | turquoise |
| ENSG00000189134 | NKAPL | 0.0022865 | yellow |
| ENSG00000129467 | ADCY4 | 0.0022664 | brown |
| ENSG00000215252 | GOLGA8B | 0.0022564 | turquoise |
| ENSG00000100439 | ABHD4 | 0.0022178 | blue |

| ensembl ID | symbol | weight | module |
|---|---|---|---|
| ENSG00000188803 | SHISA6 | 0.0021551 | blue |
| ENSG00000140577 | CRTC3 | 0.0020721 | turquoise |
| ENSG00000163046 | ANKRD30BL | 0.0020595 | yellow |
| ENSG00000228451 | SDAD1P1 | 0.002031 | grey |
| ENSG00000069122 | ADGRF5 | 0.0020031 | brown |
| ENSG00000124508 | BTN2A2 | 0.0019836 | blue |
| ENSG00000091986 | CCDC80 | 0.0019355 | blue |
| ENSG00000181026 | AEN | 0.0018551 | brown |
| ENSG00000230373 | GOLGA6L5P | 0.0018539 | yellow |
| ENSG00000216775 | AL109918.1 | 0.0017677 | turquoise |
| ENSG00000084628 | NKAIN1 | 0.0017654 | turquoise |
| ENSG00000159433 | STARD9 | 0.0016761 | turquoise |
| ENSG00000130518 | IQCN | 0.0015905 | turquoise |
| ENSG00000167771 | RCOR2 | 0.0015789 | turquoise |
| ENSG00000198804 | MT-CO1 | 0.0015404 | turquoise |
| ENSG00000144115 | THNSL2 | 0.0015386 | grey |
| ENSG00000169629 | RGPD8 | 0.0015323 | turquoise |
| ENSG00000186188 | FFAR4 | 0.0014579 | turquoise |
| ENSG00000261810 | AC133565.1 | 0.0014515 | yellow |
| ENSG00000180777 | ANKRD30B | 0.0014506 | yellow |
| ENSG00000212907 | MT-ND4L | 0.0013873 | turquoise |
| ENSG00000281708 | ERC2-IT1 | 0.0013602 | yellow |
| ENSG00000255495 | AC145124.1 | 0.0013126 | yellow |
| ENSG00000197061 | HIST1H4C | 0.0012951 | yellow |
| ENSG00000231365 | AL359915.2 | 0.0012533 | turquoise |
| ENSG00000204219 | TCEA3 | 0.0012253 | blue |
| ENSG00000179455 | MKRN3 | 0.0012236 | green |
| ENSG00000282508 | LINC01002 | 0.0011844 | turquoise |
| ENSG00000198546 | ZNF511 | 0.0011379 | grey |
| ENSG00000087086 | FTL | 0.0011253 | blue |
| ENSG00000188786 | MTF1 | 0.001069 | turquoise |
| ENSG00000144824 | PHLDB2 | 0.0010652 | turquoise |
| ENSG00000243051 | RN7SL269P | 0.001046 | yellow |
| ENSG00000136111 | TBC1D4 | 0.0010354 | turquoise |
| ENSG00000085276 | MECOM | 0.0010352 | brown |
| ENSG00000145002 | FAM86B2 | 0.00098662 | turquoise |
| ENSG00000188783 | PRELP | 0.00098385 | green |
| ENSG00000210082 | MT-RNR2 | 0.00097678 | turquoise |
| ENSG00000048540 | LMO3 | 0.0009693 | turquoise |
| ENSG00000087077 | TRIP6 | 0.00092652 | blue |
| ENSG00000123143 | PKN1 | 0.00087972 | blue |
| ENSG00000126217 | MCF2L | 0.00084918 | turquoise |
| ENSG00000254838 | GVINP1 | 0.00084161 | yellow |
| ENSG00000064703 | DDX20 | 0.00083476 | turquoise |
| ENSG00000115970 | THADA | 0.00080497 | turquoise |
| ENSG00000081479 | LRP2 | 0.00077533 | green |
| ENSG00000256193 | LINC00507 | 0.00077061 | turquoise |
| ENSG00000102057 | KCND1 | 0.00074378 | turquoise |
| ENSG00000128564 | VGF | 0.00073085 | turquoise |
| ENSG00000264920 | AC018521.5 | 0.00070618 | turquoise |
| ENSG00000145721 | LIX1 | 0.00069437 | blue |
| ENSG00000189056 | RELN | 0.00066443 | turquoise |
| ENSG00000187398 | LUZP2 | 0.00063733 | turquoise |
| ENSG00000274020 | LINC01138 | 0.00061745 | blue |
| ENSG00000274642 | AC244669.2 | 0.00061283 | turquoise |
| ENSG00000168754 | FAM178B | 0.00061088 | grey |
| ENSG00000005961 | ITGA2B | 0.00056254 | turquoise |
| ENSG00000148824 | MTG1 | 0.00054822 | turquoise |
| ENSG00000189376 | C8orf76 | 0.00054639 | turquoise |
| ENSG00000124201 | ZNFX1 | 0.00054639 | turquoise |
| ENSG00000177666 | PNPLA2 | 0.00054162 | turquoise |
| ENSG00000101187 | SLCO4A1 | 0.00053347 | brown |
| ENSG00000204685 | STARD7-AS1 | 0.00051409 | turquoise |

| ensembl ID | symbol | weight | module |
|---|---|---|---|
| ENSG00000197958 | RPL12 | 0.00050554 | turquoise |
| ENSG00000153066 | TXNDC11 | 0.00044926 | turquoise |
| ENSG00000160293 | VAV2 | 0.00044486 | turquoise |
| ENSG00000232388 | SMIM26 | 0.00044214 | turquoise |
| ENSG00000122547 | EEPD1 | 0.00043752 | turquoise |
| ENSG00000204392 | LSM2 | 0.00042365 | turquoise |
| ENSG00000177025 | C19orf18 | 0.00038519 | grey |
| ENSG00000125637 | PSD4 | 0.00038495 | blue |
| ENSG00000157399 | ARSE | 0.00038198 | blue |
| ENSG00000255182 | AC084125.2 | 0.00036771 | turquoise |
| ENSG00000171105 | INSR | 0.00036376 | green |
| ENSG00000137310 | TCF19 | 0.00035297 | brown |
| ENSG00000162623 | TYW3 | 0.0003524 | turquoise |
| ENSG00000132357 | CARD6 | 0.00034801 | brown |
| ENSG00000113721 | PDGFRB | 0.0003364 | green |
| ENSG00000087087 | SRRT | 0.00032546 | turquoise |
| ENSG00000198763 | MT-ND2 | 0.00031138 | turquoise |
| ENSG00000187867 | PALM3 | 0.00029042 | turquoise |
| ENSG00000242193 | CRYZL2P | 0.00027308 | turquoise |
| ENSG00000092295 | TGM1 | 0.00026996 | turquoise |
| ENSG00000102935 | ZNF423 | 0.00026269 | blue |
| ENSG00000250067 | YJEFN3 | 0.00025408 | turquoise |
| ENSG00000279289 | AL136164.3 | 0.00023655 | yellow |
| ENSG00000166428 | PLD4 | 0.0002263 | grey |
| ENSG00000033867 | SLC4A7 | 0.00021756 | yellow |
| ENSG00000265218 | AC103810.2 | 0.00021457 | yellow |
| ENSG00000100814 | CCNB1IP1 | 0.00021311 | turquoise |
| ENSG00000229956 | ZRANB2-AS2 | 0.00018135 | yellow |
| ENSG00000166763 | STRCP1 | 0.00017843 | turquoise |
| ENSG00000142227 | EMP3 | 0.00017697 | brown |
| ENSG00000146535 | GNA12 | 0.00017026 | blue |
| ENSG00000266644 | AC103810.5 | 0.00016552 | yellow |
| ENSG00000275620 | AL121827.2 | 0.00016073 | blue |

**Table S8 Model metrics for the linear regression between global cognitive function and the dependent variables stratified by SI modules in ROSMAP DLPFC samples. Cells with significant p values (< 0.05) are shown in bold.**

| target = global cognitive function | blue module | | | | | | brown module | | | | | | green module | | | | | |
|---|---|---|---|---|---|---|---|---|---|---|---|---|---|---|---|---|---|---|
| | Estimate | Std. Error | t value | Pr(>\|t\|) | PVE | sig. code | Estimate | Std. Error | t value | Pr(>\|t\|) | PVE | sig. code | Estimate | Std. Error | t value | Pr(>\|t\|) | PVE | sig. code |
| (Intercept) | -0.176 | 0.874 | -0.202 | 8.40E-01 | | | -0.310 | 0.898 | -0.345 | 7.30E-01 | | | -0.526 | 0.832 | -0.632 | 5.27E-01 | | |
| SI | -0.045 | 0.009 | -5.065 | **6.23E-07** | 10.45 | *** | -0.023 | 0.008 | -2.826 | **4.94E-03** | 2.34 | ** | -0.054 | 0.007 | -7.923 | **2.28E-14** | 21.10 | *** |
| age_death | -0.023 | 0.007 | -3.055 | **2.40E-03** | 3.01 | ** | -0.030 | 0.007 | -4.036 | **6.51E-05** | 5.91 | *** | -0.021 | 0.007 | -2.982 | **3.04E-03** | 2.82 | ** |
| educ | 0.034 | 0.014 | 2.468 | **1.40E-02** | 0.15 | * | 0.035 | 0.014 | 2.532 | **1.17E-02** | 0.29 | * | 0.034 | 0.013 | 2.614 | **9.29E-03** | 0.26 | ** |
| msex | -0.043 | 0.091 | -0.470 | 6.39E-01 | 0.20 | | -0.022 | 0.094 | -0.232 | 8.17E-01 | 0.12 | | -0.004 | 0.088 | -0.040 | 9.68E-01 | 0.01 | |
| race7 | -0.272 | 0.336 | -0.809 | 4.19E-01 | 0.28 | | -0.325 | 0.343 | -0.949 | 3.43E-01 | 0.32 | | -0.110 | 0.323 | -0.340 | 7.34E-01 | 0.04 | |
| apoe4 | -0.463 | 0.093 | -4.989 | **9.04E-07** | 6.05 | *** | -0.530 | 0.094 | -5.620 | **3.57E-08** | 8.19 | *** | -0.374 | 0.090 | -4.146 | **4.13E-05** | 3.86 | *** |
| RIN | 0.073 | 0.045 | 1.608 | 1.09E-01 | 0.32 | | 0.120 | 0.045 | 2.656 | **8.21E-03** | 1.14 | ** | 0.121 | 0.042 | 2.860 | **4.45E-03** | 1.23 | ** |
| PMI | 0.021 | 0.010 | 2.102 | **3.61E-02** | 0.37 | * | 0.019 | 0.010 | 1.855 | 6.43E-02 | 0.26 | . | 0.010 | 0.009 | 1.097 | 2.73E-01 | 0.02 | |
| r_pd | 0.303 | 0.058 | 5.241 | **2.58E-07** | 6.23 | *** | 0.290 | 0.059 | 4.879 | **1.54E-06** | 5.70 | *** | 0.319 | 0.055 | 5.743 | **1.83E-08** | 6.43 | *** |
| r_stroke | 0.113 | 0.065 | 1.728 | 8.47E-02 | 0.30 | . | 0.118 | 0.067 | 1.775 | 7.66E-02 | 0.33 | . | 0.099 | 0.063 | 1.578 | 1.15E-01 | 0.23 | |
| dlbdx3 | -0.756 | 0.146 | -5.188 | **3.38E-07** | 4.44 | *** | -0.749 | 0.149 | -5.028 | **7.48E-07** | 4.45 | *** | -0.557 | 0.142 | -3.924 | **1.02E-04** | 2.28 | *** |
| hspath_typ | -0.744 | 0.186 | -4.004 | **7.41E-05** | 2.62 | *** | -0.747 | 0.190 | -3.938 | **9.69E-05** | 2.64 | *** | -0.587 | 0.179 | -3.271 | **1.16E-03** | 1.60 | ** |
| arteriol_scler | -0.083 | 0.043 | -1.947 | 5.23E-02 | 0.61 | . | -0.079 | 0.044 | -1.801 | 7.25E-02 | 0.55 | . | -0.061 | 0.041 | -1.493 | 1.36E-01 | 0.33 | |
| Multiple R^2 | 0.350 | | | | | | 0.322 | | | | | | 0.402 | | | | | |
| Adjusted R^2 | 0.326 | | | | | | 0.297 | | | | | | 0.380 | | | | | |

| target = global cognitive function | grey module | | | | | | turquoise module | | | | | | yellow module | | | | | |
|---|---|---|---|---|---|---|---|---|---|---|---|---|---|---|---|---|---|---|
| | Estimate | Std. Error | t value | Pr(>\|t\|) | PVE | sig. code | Estimate | Std. Error | t value | Pr(>\|t\|) | PVE | sig. code | Estimate | Std. Error | t value | Pr(>\|t\|) | PVE | sig. code |
| (Intercept) | -1.223 | 0.903 | -1.355 | 1.76E-01 | | | 1.481 | 0.840 | 1.764 | 7.85E-02 | | . | 0.512 | 1.142 | 0.449 | 6.54E-01 | | |
| SI | 0.040 | 0.014 | 2.810 | **5.20E-03** | 3.52 | ** | -0.057 | 0.006 | -9.565 | **< 2E-16** | 23.65 | *** | -0.058 | 0.034 | -1.734 | 8.36E-02 | 2.85 | . |
| age_death | -0.032 | 0.007 | -4.399 | **1.39E-05** | 5.40 | *** | -0.020 | 0.007 | -3.008 | **2.79E-03** | 1.93 | ** | -0.029 | 0.007 | -3.932 | **9.92E-05** | 5.02 | *** |
| educ | 0.032 | 0.014 | 2.266 | **2.40E-02** | 0.05 | * | 0.036 | 0.013 | 2.869 | **4.34E-03** | 0.13 | ** | 0.031 | 0.014 | 2.208 | **2.78E-02** | 0.07 | * |
| msex | -0.063 | 0.093 | -0.671 | 5.03E-01 | 0.45 | | -0.173 | 0.086 | -2.015 | **4.46E-02** | 0.89 | * | -0.048 | 0.094 | -0.507 | 6.13E-01 | 0.22 | |
| race7 | -0.353 | 0.343 | -1.030 | 3.04E-01 | 0.18 | | -0.198 | 0.313 | -0.633 | 5.27E-01 | 0.47 | | -0.289 | 0.346 | -0.835 | 4.04E-01 | 0.30 | |
| apoe4 | -0.525 | 0.094 | -5.569 | **4.68E-08** | 7.55 | *** | -0.435 | 0.086 | -5.037 | **7.13E-07** | 4.67 | *** | -0.513 | 0.095 | -5.419 | **1.03E-07** | 7.54 | *** |
| RIN | 0.095 | 0.046 | 2.065 | **3.96E-02** | 1.31 | * | -0.144 | 0.049 | -2.911 | **3.80E-03** | 2.41 | ** | 0.096 | 0.047 | 2.041 | **4.19E-02** | 0.54 | * |
| PMI | 0.025 | 0.010 | 2.430 | 1.55E-02 | 0.27 | * | 0.015 | 0.009 | 1.598 | 1.11E-01 | 0.15 | | 0.021 | 0.010 | 2.068 | **3.93E-02** | 0.40 | * |
| r_pd | 0.289 | 0.060 | 4.858 | **1.70E-06** | 6.26 | *** | 0.272 | 0.054 | 5.037 | **7.15E-07** | 4.73 | *** | 0.309 | 0.059 | 5.200 | **3.17E-07** | 6.53 | *** |
| r_stroke | 0.123 | 0.067 | 1.849 | 6.52E-02 | 0.46 | . | 0.117 | 0.061 | 1.923 | 5.52E-02 | 0.36 | . | 0.125 | 0.067 | 1.873 | 6.18E-02 | 0.38 | . |
| dlbdx3 | -0.751 | 0.149 | -5.045 | **6.86E-07** | 4.31 | *** | -0.526 | 0.138 | -3.820 | **1.55E-04** | 2.17 | *** | -0.740 | 0.150 | -4.931 | **1.20E-06** | 4.38 | *** |
| hspath_typ | -0.780 | 0.190 | -4.103 | **4.94E-05** | 2.30 | *** | -0.630 | 0.173 | -3.630 | **3.20E-04** | 1.86 | *** | -0.734 | 0.191 | -3.841 | **1.43E-04** | 2.56 | *** |
| arteriol_scler | -0.077 | 0.044 | -1.767 | 7.81E-02 | 0.41 | . | -0.055 | 0.040 | -1.382 | 1.68E-01 | 0.27 | | -0.085 | 0.044 | -1.931 | 5.42E-02 | 0.63 | . |
| Multiple R^2 | 0.322 | | | | | | 0.437 | | | | | | 0.314 | | | | | |
| Adjusted R^2 | 0.297 | | | | | | 0.416 | | | | | | 0.289 | | | | | |

Signif. codes: 0 ‘***’ 0.001 ‘**’ 0.01 ‘*’ 0.05 ‘.’ 0.1 ‘ ’ 1
