## supplemental materials for "Deep learning-based brain transcriptomic signatures associated with the neuropathological and clinical severity of Alzheimer’s disease"

**Supplementary material**

**Supplementary Figure legends**

**Figure S1 Hyper-parameter estimation and model training. a)** The regularization parameter λ was estimated to be 0.004 based on the one-standard-error rule. **b)** The trade-off parameter α was estimated to be 2. **c)** Curves of training and validation losses vs. epochs. No sign of over-fitting was observed. The first 1500 epochs are for model initialization based on supervised learning and the remaining 5000 epochs are for model optimization. **d)** Curves of training and validation accuracy vs. epochs.

**Figure S2 Index gene identification.** **a)** Histogram of the weights in logarithm scale for each gene in the training model. **b)** Volcano plot showing the distribution of the index genes vs all the other genes in ROSMAP dataset. P and log_2_FC values were obtained from DEG analysis (syn8456629). **c)** Volcano plot showing the distribution of the index genes vs all the DEGs identified in the meta-analysis of AMP-AD datasets (syn11914606). P and log_2_FC values were the same as b), taken from ROSMAP dataset (syn8456629), but genes not identified as DEGs in the meta-analysis are not shown for clarity.

**Figure S3 Regression plots between SI and global cognitive function for the index genes from each module. a)** blue; **b)** brown; **c)** green; **d)** grey; **e)** turquoise; **f)** yellow.

**Figure S4 Regression plots between SI and global cognitive function for the stratified groups by diagnosis.** **a)** All AD and control subjects (used in supervised model training); **b)** All other subjects (not used in supervised model training). The fitting lines for all subjects (bold) and stratified group (grey) are also shown.

**Figure S5 The SI modules were examined for overlap (Fisher’s exact test) with the curated AD gene sets and co-expression modules derived from the individual dataset of AMP-AD cohorts.** Overlaps were shown for those adjusted p (FDR correction) < 0.05. Correction was done independently for each set or each study. **a)** The curated gene sets were taken from Genecards,^1^ GeneRIF,^2^ Panther,^3^ dbGaP,^4^ IGAP,^5^ OMIM,^6^ KEGG,^7^ and WikiPathway^8^ respectively. The module genes were taken from **b)** Mayo network^9^, **c)** MSBB network^10^, **d)** ROSMAP network^11^. Only those modules reported as associated and/or top ranked with AD traits were included in Figure **b)-d)**.

**Figure S6 Comparisons between SI and pseudotime from the work of Mukherjee et al. obtained on the same subjects (n = 537) in the ROSMAP cohort. a)** Correlation plots between SI and pseudotime in female and male samples respectively. Correlation coefficients were obtained by lm function in R, with no covariates included. **b)** Distribution of samples’ SI for three different external measures of AD staging: Braak score (tau pathology), CERAD score (amyloid pathology), and cognitive diagnosis (clinical measure of disease severity), stratified by sex (in comparison with Figure 3 and S9 in Mukherjee et al). **c)** Association p-values between SI/pseudotime and the three external measures, by linear or logistic regression, stratified by sex.

**Figure S7 Cell-type gene expression signatures as a function of SI. Mean expression of cell markers for astrocytes, neurons, microglia, oligodendrocytes and endothelial cells were plotted and colored respectively. a)** all subjects. **b)** female only. **c)** male only.

**Supplementary Table legends**

**Table S1 Demographic information for all the subjects included in RNA-seq data used in this study.**

**Table S2 Description of the variables used in the linear regression model for ROSMAP cohort.**

**Table S3 Model metrics for the linear regression between global cognitive function and the dependent variables (as defined in Table S2) stratified by diagnosis groups in ROSMAP DLPFC samples.**

**Table S4 Model metrics (p values and correlation coefficients) for the linear regression between all the neuropathological biomarkers and the dependent variables (as defined in Table S2) stratified by diagnosis groups in ROSMAP DLPFC samples.**

**Table S5 Model metrics (p values and correlation coefficients) for the linear regression between neuropathological biomarkers and the dependent variables for the two brain regions in Mayo samples.**

**Table S6 Model metrics (p values and correlation coefficients) for the linear regression between neuropathological and clinical biomarkers and the dependent variables for the four brain regions in MSBB samples.**

**Table S7 Index genes and their weights contributing to the deep learning model.**

**Table S8 Model metrics for the linear regression between global cognitive function and the dependent variables stratified by SI modules in ROSMAP DLPFC samples.**
